# Progenitor identification and SARS-CoV-2 infection in long-term human distal lung organoid cultures

**DOI:** 10.1101/2020.07.27.212076

**Authors:** Ameen A. Salahudeen, Shannon S. Choi, Arjun Rustagi, Junjie Zhu, Sean M. de la O, Ryan A. Flynn, Mar Margalef-Català, António J. M. Santos, Jihang Ju, Arpit Batish, Vincent van Unen, Tatsuya Usui, Grace X.Y. Zheng, Caitlin E. Edwards, Lisa E. Wagar, Vincent Luca, Benedict Anchang, Monica Nagendran, Khanh Nguyen, Daniel J. Hart, Jessica M. Terry, Phillip Belgrader, Solongo B. Ziraldo, Tarjei S. Mikkelsen, Pehr B. Harbury, Jeffrey S. Glenn, K. Christopher Garcia, Mark M. Davis, Ralph S. Baric, Chiara Sabatti, Manuel R. Amieva, Catherine A. Blish, Tushar J. Desai, Calvin J. Kuo

## Abstract

The distal lung contains terminal bronchioles and alveoli that facilitate gas exchange and is affected by disorders including interstitial lung disease, cancer, and SARS-CoV-2-associated COVID-19 pneumonia. Investigations of these localized pathologies have been hindered by a lack of 3D *in vitro* human distal lung culture systems. Further, human distal lung stem cell identification has been impaired by quiescence, anatomic divergence from mouse and lack of lineage tracing and clonogenic culture. Here, we developed robust feeder-free, chemically-defined culture of distal human lung progenitors as organoids derived clonally from single adult human alveolar epithelial type II (AT2) or KRT5^+^ basal cells. AT2 organoids exhibited AT1 transdifferentiation potential, while basal cell organoids progressively developed lumens lined by differentiated club and ciliated cells. Organoids consisting solely of club cells were not observed. Upon single cell RNA-sequencing (scRNA-seq), alveolar organoids were composed of proliferative AT2 cells; however, basal organoid *KRT5*^+^ cells contained a distinct *ITGA6*^+^*ITGB4*^+^ mitotic population whose proliferation segregated to a *TNFRSF12A^hi^* subfraction. Clonogenic organoid growth was markedly enriched within the TNFRSF12A^hi^ subset of FACS-purified ITGA6^+^ITGB4^+^ basal cells from human lung or derivative organoids. *In vivo*, TNFRSF12A^+^ cells comprised ~10% of KRT5^+^ basal cells and resided in clusters within terminal bronchioles. To model COVID-19 distal lung disease, we everted the polarity of basal and alveolar organoids to rapidly relocate differentiated club and ciliated cells from the organoid lumen to the exterior surface, thus displaying the SARS-CoV-2 receptor ACE2 on the outwardly-facing apical aspect. Accordingly, basal and AT2 “apical-out” organoids were infected by SARS-CoV-2, identifying club cells as a novel target population. This long-term, feeder-free organoid culture of human distal lung alveolar and basal stem cells, coupled with single cell analysis, identifies unsuspected basal cell functional heterogeneity and exemplifies progenitor identification within a slowly proliferating human tissue. Further, our studies establish a facile *in vitro* organoid model for human distal lung infectious diseases including COVID-19-associated pneumonia.

## INTRODUCTION

The distal lung, including terminal bronchioles and alveoli, performs essential gas exchange functions which can be significantly compromised by disease. For example, SARS-CoV-2 infection can elicit severe distal lung COVID-19 pathology with life-threatening pneumonia and respiratory failure^1–3^. The limited understanding of COVID-19 pneumonia pathogenesis during a worldwide pandemic has highlighted a pressing need for robust *in vitro* culture systems allowing study of distal lung pathologies in primary human cells.

The lack of long-term human distal lung culture systems has precluded functional testing of the proliferative capacity of putative human distal lung stem cell populations, which are therefore largely inferred from mouse studies. In mouse, lineage-tracing has enabled *in vivo* confirmation and mapping of ‘bifunctional’ distal lung stem cells that constitutively execute both physiologic and regenerative functions, namely secretory club cells in distal bronchioles^4,5^ and surfactant-producing alveolar epithelial type II (AT2) cells in alveoli^6,7^. Injury-inducible murine lung populations include an alveolar progenitor that renews AT1 and AT2 cells^8^, and distal airway basal cell-like^9–11^ or bronchioalveolar^12,13^ progenitors with airway and alveolar differentiation potential. Whether the human correlates of these mouse stem cells are functional in renewing mature lung cell lineages is largely unknown.

The cell type composition of human terminal airways differs substantially from mouse. In the human lung basal cells span the entire airway axis^14^, while in mouse they are absent from the terminal bronchioles where club cells renew and repair the epithelium^15,16^. Long-term culture of human tracheal and bronchial basal cells have demonstrated stem cell potential, and these are also presumed to function as stem cells for lower airway renewal^17–23^. Human AT2 cell cultures have also been reported but are short-lived^6,18,24,25^, and the long-term self-renewal capacity of human AT2 cells thus remains unknown. Furthermore, existing AT2 cell culture protocols achieve only minimal expansion and require feeder cells producing unknown factors, limiting their biological characterization and screening utility. Directed differentiation of induced pluripotent stem cells (iPSCs) to AT2 cells can be limited by efficiency, feeder dependence and persistent fetal gene expression, suggesting immaturity^26–29^. Here, we established long-term, feeder-free, chemically-defined 3D organoid culture of distal human lung, including AT2 and basal stem cells, and applied this method to progenitor identification and SARS-CoV-2 modeling.

### Human adult distal lung culture yields clonogenic alveolar and basal cell organoids

We empirically established defined media conditions supporting clonal expansion of distal human lung progenitors, encompassing bronchiolar and alveolar cells. To exclude proximal cell types, we used only the peripheral one centimeter of lung underlying the mesothelium, which was devoid of cartilage **(Fig. 1a)**. Single cell suspensions from 134 individuals **(Supplementary Table 1)**, were cultured within a droplet of collagen/laminin extracellular matrix without exogenous feeder cells. We surveyed growth factors whose cognate pathways including WNT, EGF and BMP have been implicated in lung development and disease pathogenesis^14,22^. The combination of EGF and the BMP antagonist NOGGIN was optimal, without any additional growth-promoting effects of either WNT3A or R-SPONDIN1 (RSPO1) **(Extended Data Fig. 1a,b)**. Single cells underwent clonal expansion into one of two distinct organoid morphologies. Cystic organoids **(Fig. 1a,b)** were SFTPC^+^HT2-280^+^ and lacked KRT5, indicating AT2 cell identity (**Fig. 1d-g**). In contrast, solid organoids **(Fig 1a,c)** expressed the basal cell marker KRT5 and lacked SFPTC indicating basal cell identity **(Fig. 1d, h-j)**. Organoids consisting solely of SCGB1A1^+^ club cells were not observed **(Fig. 1d)**.

**Figure 1.**
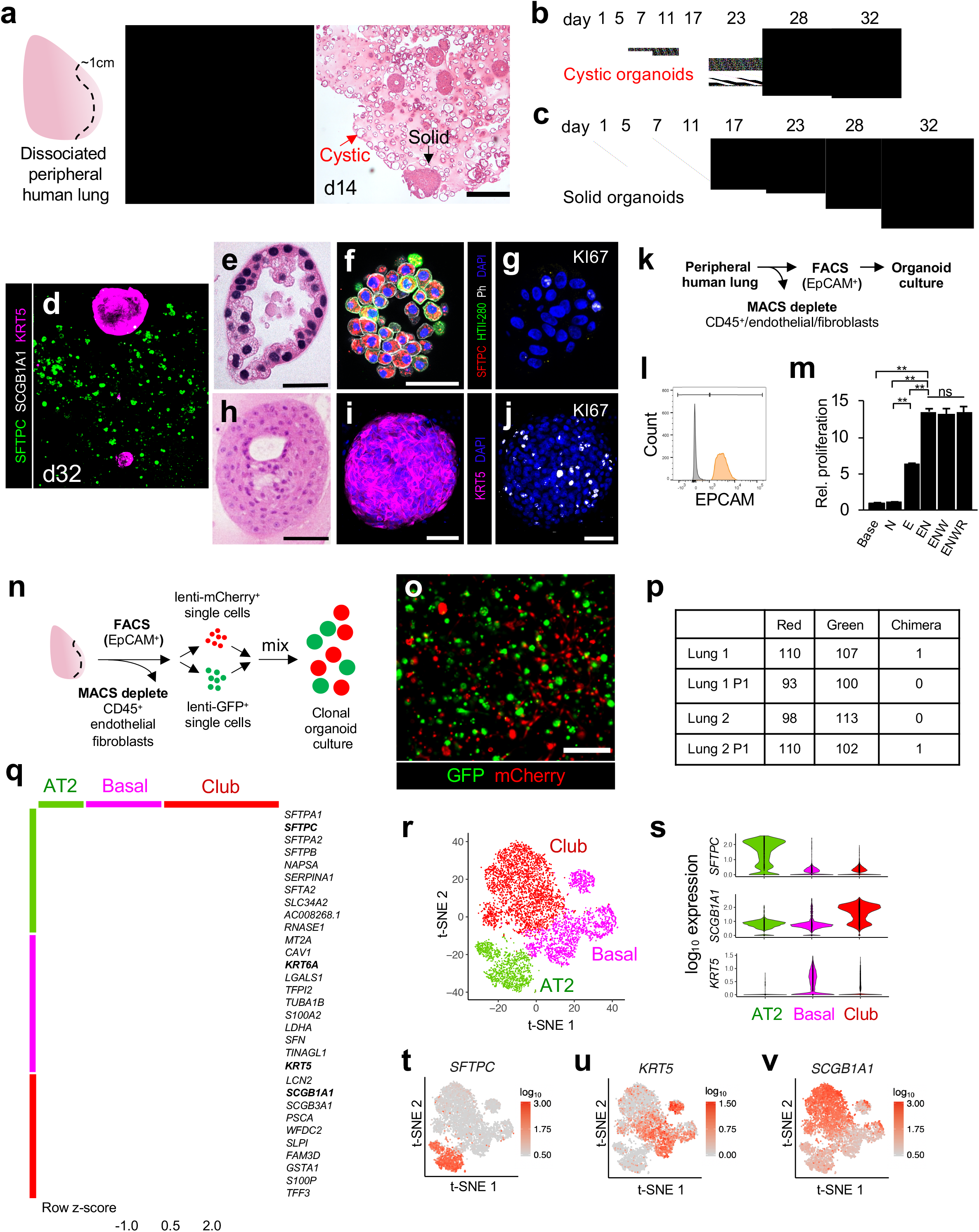
Clonogenic expansion of human distal lung organoids in chemically defined conditions. **a.** Day 14 organoid culture of dissociated unfractionated human distal lung, H&E. Cystic and solid organoids are denoted. Scale bar = 100 μm. **b-c,** Time lapse transmission confocal images of solid (b) and cystic (c) organoids originating from single cells, scale bar = 100 μm. **d,** Combinatorial whole-mount immunofluorescence (IF) of organoid cultures for anti-KRT5 (basal) SCGB1A1 (club) and SFTPC (AT2), scale bar = 100 μm, day 32. **e-g,** Analysis of alveolar organoids, day 32. **e,** H&E image of cystic AT2 organoid, scale bar = 25 μm. **f,** Combinatorial whole-mount fluorescence for anti-SFTPC, anti HTII-28, phalloidin and DAPI, scale bar = 50 μm. **g,** anti-Ki67 and DAPI fluorescence of adjacent section of (e) **h-j,** Analysis of basal organoids, day 32. **h,** H&E of basal organoid, scale bar = 50 μm. **i,** Combinatorial whole-mount fluorescence for anti-KRT5 and DAPI, scale bar =100 μm**, j,** Ki67 immunostaining of (i). **k,** Purification schema to isolate epithelial cells from distal human lung involving negative MACS bead depletion of CD45^+^ hematopoietic cells, endothelial cells and fibroblasts, followed by positive FACS selection for EPCAM^+^ epithelium. **l,** Representative FACS from the purification of (k) demonstrating > 99.9% EPCAM^+^ purity (orange) upon re-analysis versus unstained controls (grey). **m,** Proliferation of EPCAM^+^ cells purified from distal lung as in (k) after day 10 of organoid culture with specified growth factors N=Noggin, E=EGF, W=WNT3A, R=RSPO1 from 3 technical replicates, error bars = SEM, *= p < .05 **n-p,** Clonality mixing studies. **n,** Schema of mixing studies of lentivirus-GFP- and lentivirus-mCherry-expressing cells to determine clonality. **o,** Representative live fluorescent imaging of resultant green and red organoids from (m), scale bar = 500 μm. **p,** Quantitation of red, green, or chimeric, distal lung organoid cultures from two separate lung donors (1, 2) after initial and serial passaging (P1=passage 1). **q-v**, scRNA-seq of day 28 total distal lung organoid cultures. **q,** Unsupervised clustering of scRNA-seq of day 28 total distal lung organoid cultures demonstrates AT2, basal, and club cell populations, with canonical markers of these cell types (*SFTPC* (AT2), *KRT6A/KRT5* (Basal), *SCGB1A1* (club)). **r,** t-SNE plot of 7,285 individual cells from (q) displays the cell classes. **s,** Violin plots of (r). **t-v,** Feature plots highlight distribution and log10 UMI counts corresponding to q-s.* p < 0.05, ** p < 0.01, two-tailed Student’s t-test; ns, non-significant. For scRNA-seq analysis in q-v, a modified Kruskal-Wallis Rank Sum Test was performed to determine significance of differential marker gene expression for AT2, basal, and club, with all p-values < 0.001.

To stringently exclude the possibility that contaminating stromal cells could be contributing unknown growth factors, we generated distal lung organoids from >99.9% EPCAM^+^ starting populations by magnetic bead depletion of fibroblasts, endothelial and hematopoietic cells followed by FACS purification of EPCAM^+^ cells, which confirmed organoid generation with only EGF and NOGGIN provision (**Fig. 1k-m**). Distal lung organoids could be passaged for ~6 months with basal organoids initially exhibiting 6-7 doublings every 2 weeks. Alveolar organoids expanded more slowly with an initial rate of 3-4 doublings/2 weeks but predominated over basal organoids after several months. Based on initial cell division rates, the upper limits of basal and alveolar expansion were 2^19^ (524,288 fold) and 2^16^ (65,536 fold) respectively. Organoids arose clonally as confirmed by (1) time-lapse microscopy of single cells (**Fig. 1b,c**) and (2) color mixing studies of disaggregated, lentivirally-transduced GFP^+^ or mCherry^+^ cells that generated entirely red or green but not chimeric organoids **(Fig. 1n-p)**.

Single cell RNA-sequencing of distal lung organoids confirmed distinct *SFTPC*^+^ AT2, *KRT5*^+^ basal and *SCGB1A1*^+^ club cell populations. Cells co-expressing *KRT5* and *SCGB1A1* bridged the basal and club cell populations, suggesting molecular intermediates transitioning from basal into club cells (**Fig. 1q-v, Extended Data Fig. 2**). Trajectory analysis using SPADE^30,31^ projected a cellular differentiation path from basal to club cell identity but not between AT2 and club or AT2 and basal cells (**Extended Data Fig. 3, Supplementary Data 1**).

### Human AT2 cells extensively renew and maintain AT1 cell transdifferentiation capacity

We further refined these methods to generate pure clonogenically-derived AT2 organoids. Viable AT2 cells were purified from mixed distal lung organoids without accompanying stromal populations using fluorescence-associated cell sorting (FACS), exploiting lamellar body uptake of the lysosomal dye LysoTracker^32^ in EPCAM^+^ cells **(Fig. 2a**, **Extended Data Fig. 1c-d, Extended Data Fig. 4a, Supplementary Data 2).** Individual EPCAM^+^LysoTracker^+^ AT2 cells progressively expanded as cystic organoids up to 180 days, exhibiting a mixture of cuboidal or more flattened morphologies reminiscent of alveoli **(Fig. 2b,c)**. Qualitatively identical results were obtained with anti-HTII-280 (AT2 marker) purification instead of LysoTracker. Transmission electron microscopy (TEM) revealed a basal surface contacting surrounding matrix, characteristic apical microvilli, and abundant cytoplasmic lamellar bodies, characteristic of fully mature and functional AT2 cells **(Fig. 2d, Extended Data Fig. 4b)**. AT2 organoids expressed HTII-280 **(Fig. 2e)** and occasionally assumed an AT1-like flattened morphology with variable downregulation of AT2 cell type markers but without AT1 marker expression (data not shown). However, upon culture on a glass surface with fetal bovine serum which promotes the AT1 phenotype^7,33^, AT2 cells rapidly flattened, downregulated HTII-280 and initiated AT1 HTI-56 marker expression, indicating retention of differentiation capacity after expansion **(Fig. 2f)**. EGF and NOGGIN were sufficient for clonal AT2 organoid proliferation and exogenous WNT-3A and RSPONDIN-1 did not enhance growth. However, the PORCUPINE inhibitor C59, which blocks endogenous WNT biosynthesis, attenuated AT2 organoid growth **(Fig. 2g),** suggesting essential autocrine WNT signaling and recapitulating mouse AT2 cell biology^34^. Lastly, isolated AT2 cells exhibited clonal organoid growth in lentivirus GFP/mCherry mixing studies, with 0/797 organoids demonstrating chimerism **(Fig. 2h)**.

**Figure 2.**
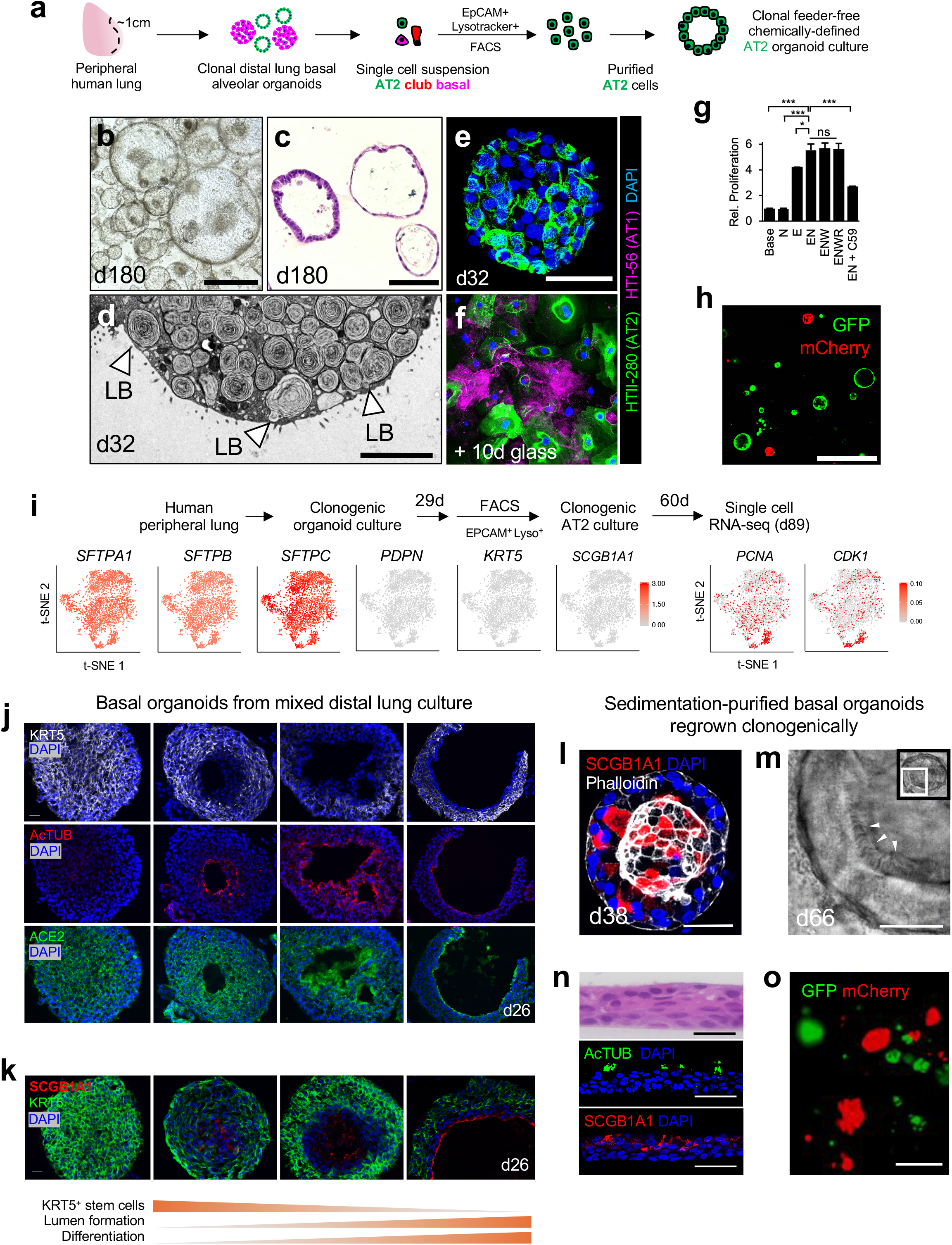
Long-term clonogenic culture of human basal and AT2 cells. **a,** Schema of FACS isolation of AT2 cells from human mixed distal lung organoids as EPCAM^+^LysoTracker^+^ AT2 cells followed by long-term clonogenic culture. **b,** AT2 organoid culture from (a), Brightfield day 180, scale bar = 200 μm. **c,** H&E from (b), day 180, scale bar = 50 μm**. d,** Transmission electron microscopy image of representative AT2 organoid from b-c, day 32, LB = lamellar body, scale bar = 5 μm. **e,** Combinatorial IF staining for AT1 (HTI-56) and AT2 (HTII-280) cell markers in AT2 organoids as in (a), day 32, scale bar = 50 μm. **f,** AT2 organoids from (e) dissociated into single cells and cultured on glass with DMEM/F12 and 5% fetal calf serum for 10 days, scale bar = 50 μm. **g,** AT2 organoid proliferation with differing combinations of niche factors and PORCUPINE inhibitor C59 (1 μM) from 3 technical replicates, error bars = SEM, * = p < 0.05 **h,** Representative image of clonal mixing studies from stroma-depleted, EPCAM^+^Lysotracker^+^ and lentivirally marked AT2 cells demonstrating presence of mCherry^+^ or GFP^+^ but not chimeric organoids carried out as in (Fig. 1n), passage 1 after lentiviral infection, scale bar = 200 μm. **i,** scRNA-seq analysis of clonal AT2 organoids from EPCAM^+^Lysotracker^+^ cells highlighting homogeneous distribution of AT2 *SFTPA1/B/C* as well as mitotic (*PCNA*, *CDK1*) mRNAs but absence of AT1 (*PDPN*), basal (*KRT5*) and club (*SCGB1A1*) mRNAs among 2,780 single cells from enriched AT2 organoids. log10 UMI counts, day 89 cumulative culture. **j-k,** Spontaneous lumen formation and differentiation of basal organoids in day 26 mixed distal lung culture. A spectrum of formation of interior lumens lined by acetylated tubulin^+^ (AcTUB) ciliated and SCGB1A1^+^ club cells, along with the SARS-CoV-2 receptor ACE2 is observed, scale bar = 20 μm. **l-o**, Clonogenic culture of human distal airway basal organoids. **l,** Spontaneous club cell differentiation within basal culture, day 38, scale bar = 50 μm. **m,** Ciliated organoid confocal transmission image from **Supplementary Video 1**, scale bar = 20 μm. **n**, H&E and immunostaining of ciliated and club cell markers of 2D air liquid interface cultures initiated from d14 Ficoll-sedimented basal cell organoids, scale bar = 50 μm. **o,** Representative image of clonal mixing studies from stroma-depleted, Ficoll-purified and lentivirally-marked basal organoid cells demonstrating mCherry^+^ or GFP^+^ but not chimeric organoids as in Fig. 1n, passage 1 after lentiviral infection, scale bar = 200 μm.

Single cell RNA-seq of mixed distal lung organoids revealed uniformly high-level expression of canonical AT2 cell markers such as *SFTPC* within the alveolar populations (**Fig. 1q-t, Extended Data Fig. 2**). These data did not readily identify AT2 cell subsets within organoids, although the relatively low number of AT2 cells limited sensitivity **(Extended Data Fig. 2, Extended Data Fig. 5a-c)**. We thus generated clonally derived pure alveolar organoids by culturing FACS-isolated EPCAM^+^Lysotracker^+^ cells from mixed distal lung organoids **(Fig. 2i)**. Analysis of 2,780 single cells from purified alveolar organoids at culture day 89 (60 days post-FACS purification) re-demonstrated homogeneous AT2 marker expression including *SFTPA1, SFTPB* and *SFTPC* without basal, club, or AT1 markers **(Fig. 2i, Supplementary Data 3-4)**. Importantly, cell cycle mRNAs such as *PCNA* and *CDK1* were expressed by a minority of AT2 cells but did not cluster to a specific population **(Fig. 2i)** and a proliferative sub-cluster within AT2 organoids having distinct gene expression unrelated to cell cycle status was not detected **(Extended Data Fig. 5d-h, Supplementary Data 4)**.

### Spontaneous differentiation of human distal airway basal cell-derived organoids

Basal cell organoids in mixed distal lung culture initially formed solid KRT5^+^ masses **(Fig. 1a, c-d).** However, by ~ 1 month, approximately 50% stochastically developed single but occasionally multiple lumens; this lumen formation did not result from apoptosis **(Fig. 2j, Extended Data Fig. 1e-g).** Lumen appearance coincided strongly with the emergence of differentiated acetylated tubulin^+^ (AcTUB^+^) ciliated cells and SCGB1A1^+^ club cells at the lumenal surface. Conversely, the basal stem cell marker KRT5 was specifically excluded from the differentiated lumen zone, but was otherwise diffusely expressed **(Fig. 2j,k)**.

We further established pure basal cell cultures by using density sedimentation to remove cystic AT2 organoids, leaving behind solid basal organoids which were then disaggregated and regrown from single cell suspensions. After 2-4 weeks in culture post-sedimentation, proliferating basal cell organoids again progressively cavitated **(Extended Data Fig. 1h-i)** with appearance of luminal SCGB1A1^+^ club and AcTUB^+^ ciliated cells either upon organoid culture **(Fig. 2l,m, Supplementary Video 1-2)** or conversion of 3D organoids to 2D air-liquid interface monolayers **(Fig. 2n)**. Clonal outgrowth of basal cell-derived organoids was confirmed by density sedimentation followed by FACS isolation and culture of EPCAM^+^ cells and culture which exhibited monoclonality in lentivirus GFP/mCherry mixing studies with only 2/845 chimeric organoids **(Fig. 2o).** Under these conditions EGF and Noggin were again sufficient for maximal growth without additive effects of WNT3A, R-SPONDIN1. Unlike AT2 organoids, growth was not significantly inhibited by the PORCUPINE inhibitor C59 **(Extended Data Fig. 1j).**

### Organoid scRNA-seq reveals two molecularly distinct subtypes of human distal airway basal cells

In contrast to the homogeneity of organoid AT2 cells, scRNA-seq clustering of *KRT5*^+^ basal cells from multiple individuals reproducibly identified two subpopulations, designated Basal 1 and Basal 2 **(Fig 3a-b, Extended Data Fig. 6).** Basal 1 was enriched for differentiation and cell fate determinants such as *HES1* and *ID1* and included an actively cycling subpopulation expressing proliferation markers *PCNA* and *CDK1* with significantly overrepresented GSEA cell cycle processes **(Fig. 3a, Extended Data Fig. 7a-b, Supplementary Data 4)**. Basal 1 but not Basal 2 included canonical lung basal cell mRNAs such as integrin α_6_ (*ITGA6*) and *TP63*^16^, as well as integrin β_4_ (*ITGB4*) which is a binding partner for integrin α_6_^35^ and also expressed in murine Lineage Negative Epithelial Progenitors (LNEPs)^9^ **(Fig. 3c)**. Basal 2, while lacking the above Basal 1 proliferative and cell fate markers, was enriched in vesicular transport, endoplasmic reticulum housekeeping processes and squamous markers; these transcripts were also present in Basal 1, albeit at lower levels **(Supplementary Data 4)**.

**Figure 3.**
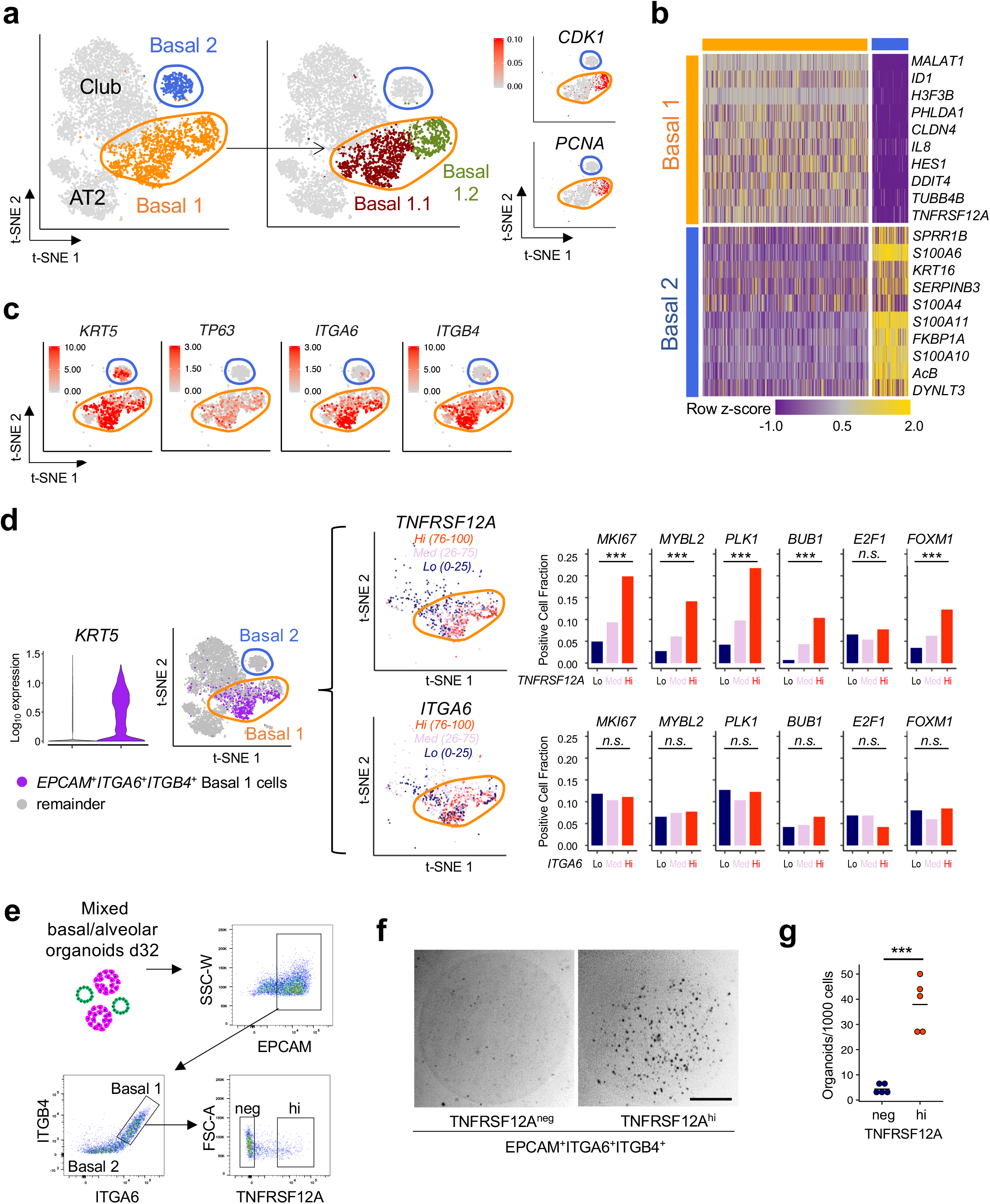
scRNA-seq-based discovery and functional validation of a proliferative human TNFRSF12A^hi^ basal cell subset in organoids. **a-c,** scRNA-seq analysis and subclustering of *KRT5*^+^ basal cells from distal lung organoid culture day 28. **a,** Basal cells from Fig. 1r are subclustered into Basal 1 (orange), characterized by proliferation and developmental programming and Basal 2 (blue), enriched for structural, cytoskeletal and calcium binding protein gene expression, t-SNE. **b,** Heat map of (a). **c,** Feature plots of (b) highlight selective enrichment of basal marker transcripts in Basal 1 versus Basal 2. log10 UMI counts are indicated. **d,** Left, Violin plot of scRNA-seq analysis from Fig. 4a depicting *KRT5* expression among *EPCAM*^+^*ITGA6*^+^*ITGB4*^+^ single cells (purple, i.e. tandem expression of all three genes) versus the remainder of cells (gray), p < 0.001 Kruskal-Wallis Rank Sum Test. Middle, t-SNE visualization of *TNFRSF12A* and *ITGA6* expression from the left panel among cells with *EPCAM*^+^*ITGA6*^+^*ITGB4*^+^ gene expression and subdivision by high (top quartile, orange), medium (pink) and low (bottom quartile, navy blue) mRNA expression. Right, Proliferation-associated gene expression is progressively enriched for scRNA-seq cell fractions of in *EPCAM*^+^*ITGA6*^+^*ITGB4*^+^ cells that are stratified for low, medium, or high expression of *TNFRSF12A* mRNA but for similar gradations of *ITGA6* mRNA, n.s. = not significant, *** = p < 0.001 Chi-square test. **e-g,** Prospective isolation and clonogenicity of TNFRSF12A^hi^ cells from mixed distal lung organoids. **e,** FACS gating strategy to subfractionate Basal 1 into TNFRSF12A^hi^ versus TNFRSF12A^neg^ from (EPCAM^+^ITGA6^+^ITGB4^+^) Basal 1 cells. Populations shown were pre-gated on live singlets. **f,** Representative brightfield image of TNFRSF12A^hi^ versus TNFRSF12A^neg^ fractions from (f) after 14 days of organoid culture. **g,** Quantitation of organoid formation in (f), each data point represents the mean of technical replicates of an organoid culture from a unique individual, *** = p < 0.001 two-tailed Student’s t-test.

### TNFRSF12A marks an organoid basal cell subpopulation with enriched progenitor activity

We next examined the membrane receptor *TNFRSF12A* (Fn14, TweakR), one of the most differentially expressed genes in the Basal 1 cluster **(Fig. 3b, Supplementary Table 2)**, because of its potential utility for FACS sorting and since a related TNF superfamily membrane receptor gene family member, *TNFRSF19*, marks gastric and intestinal stem cells^36,37^. Indeed, *TNFRSF12A* mRNA was enriched in *EPCAM*^+^*ITGA6*^+^*ITGB4*^+^ Basal 1 versus Basal 2 cells **(Fig. 3d)**. In scRNA-seq, Basal 1 *EPCAM*^+^*ITGA6*^+^*ITGB4*^+^ cells could be divided into *TNFRSF12A*-low, -medium and -high mRNA-expressing fractions, and notably a proliferative gene module^38^ was significantly enriched in the highest (*TNFRSF12A^hi^*) versus lowest quartile (*TNFRSF12A^lo^*) subsets **(Fig. 3d)**. Crucially, low, medium and high expression of the Basal 1 transcripts *ITGA6* or *TP63*, or *KRT5* did not enrich for this proliferative signature, indicating particular discriminatory utility of *TNFRSF12A* mRNA levels **(Fig. 3d, Extended Data Fig. 7c)**. To functionally validate this observation, we fractionated total distal lung organoids by anti-TNFRSF12A monoclonal antibody FACS into EPCAM^+^ITGA6^+^ITGB4^+^ cells and then into TNFRSF12A^hi^ and TNFRSF12A^neg^ subsets. The incidence of TNFRSF12A^hi^ cells declined with time to comprise a minority of EPCAM^+^ITGA6^+^ITGB4^+^ cells in late stage differentiating cultures **(Fig. 3e)**. Notably, organoid-derived TNFRSF12A^hi^ basal cells reproducibly exhibited 4-12x greater clonogenic organoid-forming capacity than TNFRSF12A^neg^ cells in 5 out of 5 individuals **(Fig. 3f,g)** in parallel with >60-fold enrichment of TNFRSF12A mRNA in the former (**Supplementary Data 2**).

Possible lineage relationships between Basal 1 and Basal 2 were examined by FACS purification followed by clonogenic culture. Density sedimentation-purified KRT5^+^ basal organoids **(Fig. 2l-n)** were fractionated into EPCAM^+^ITGA6^+^ITGB4^+^TNFRSF12A^hi^ (Basal 1) and EPCAM^+^ITGA6^-^ITGB4^-^ TNFRSF12A^neg^ (Basal 2) populations **(Extended Data Fig. 8a)**. Despite essentially homogeneous KRT5 expression in both fractions **(Extended Data Fig. 8b**) clonogenic organoid formation was strongly enriched in Basal 1 versus Basal 2 from three separate individuals **(Extended Data Fig. 8c-d)**, indicating the lack of Basal 1 cell generation by Basal 2 cells. Basal 2-enriched loci *SPRR1B* and *TMSB4X* **(Fig. 3b, Supplementary Data 4)** were transiently induced in organoids from FACS-isolated TNFRSF12A^hi^ Basal 1 cells (**Extended Data Fig. 8e-f),** indicating Basal 2 differentiation from Basal 1. SPADE trajectory analysis also demonstrated Basal 2 cells emanating from TNFRSF12A^+^ basal cells, independently supporting a one-way lineage route from Basal 1 to Basal 2 **(Extended Data Fig. 3)**.

The NOTCH target gene *HES1* was one of the most differentially upregulated Basal 1 genes, and Basal 1-upregulated gene networks included *NOTCH1, NOTCH2* and *JAG1* **(Fig. 3b, Supplementary Data 4)**. In basal organoids from FACS-purified TNFRSF12A^hi^EPCAM^+^ITGA6^+^ITGB4^+^ cells, NOTCH inhibition by the gamma-secretase inhibitor DBZ or the extracellular domain of the DLL4 E12 mutant^39^ significantly increased proliferation **(Extended Data Fig. 8g-h)**, suggesting NOTCH restrains growth. NOTCH inhibition by DBZ or DLL4 E12 NOTCH signaling also stimulated incomplete alveolar TNFRSF12A^hi^ differentiation by upregulating *SFTPC* mRNA without lamellar body or SFTPC protein production (data not shown), mirroring the effect of Notch on mouse LNEP stem cells^9^. Conversely, NOTCH agonism by JAG1 peptide did not affect proliferation but induced *SCGB1A1*, similar to reports in upper airway cells^40–42^ **(Extended Data Fig. 8i)**.

### TNFRSF12A^+^ basal cells cluster within distal airways in vivo and exhibit enhanced clonogenic potential

To demonstrate the TNFRSF12A-expressing basal subpopulation *in vivo* we performed TNFRSF12A antibody^43^ staining of freshly fixed intact human distal lung specimens. KRT5 and/or p63 marked essentially all distal airway basal cells, but TNFRSF12A strikingly labeled a cell subset localized in sporadic clusters within the basal layer **(Fig. 4a-c).** KRT5^+^TNFRSF12A^+^ basal cells frequently, but not exclusively, resided at tips or bases of bronchiolar furrows, the latter a previously recognized niche for goblet cells^44^. TNFRSF12A was not restricted to the basal layer and was detected in diverse lung stromal and epithelial cells, yet clearly marked a minor population of KRT5^+^/p63^+^ basal cells **(Fig. 4a-c)**. The TNFRSF12A-expressing subset of KRT5^+^ basal cells exhibited a higher mitotic index than total KRT5^+^ cells *in vivo*, consistent with enriched proliferative capacity *in vitro* **(Fig. 4d-e)**. FACS analysis of freshly dissociated human lung confirmed TNFRSF12A expression in 10.9% of KRT5^+^ basal cells **(Fig. 4f, top)**.

**Figure 4.**
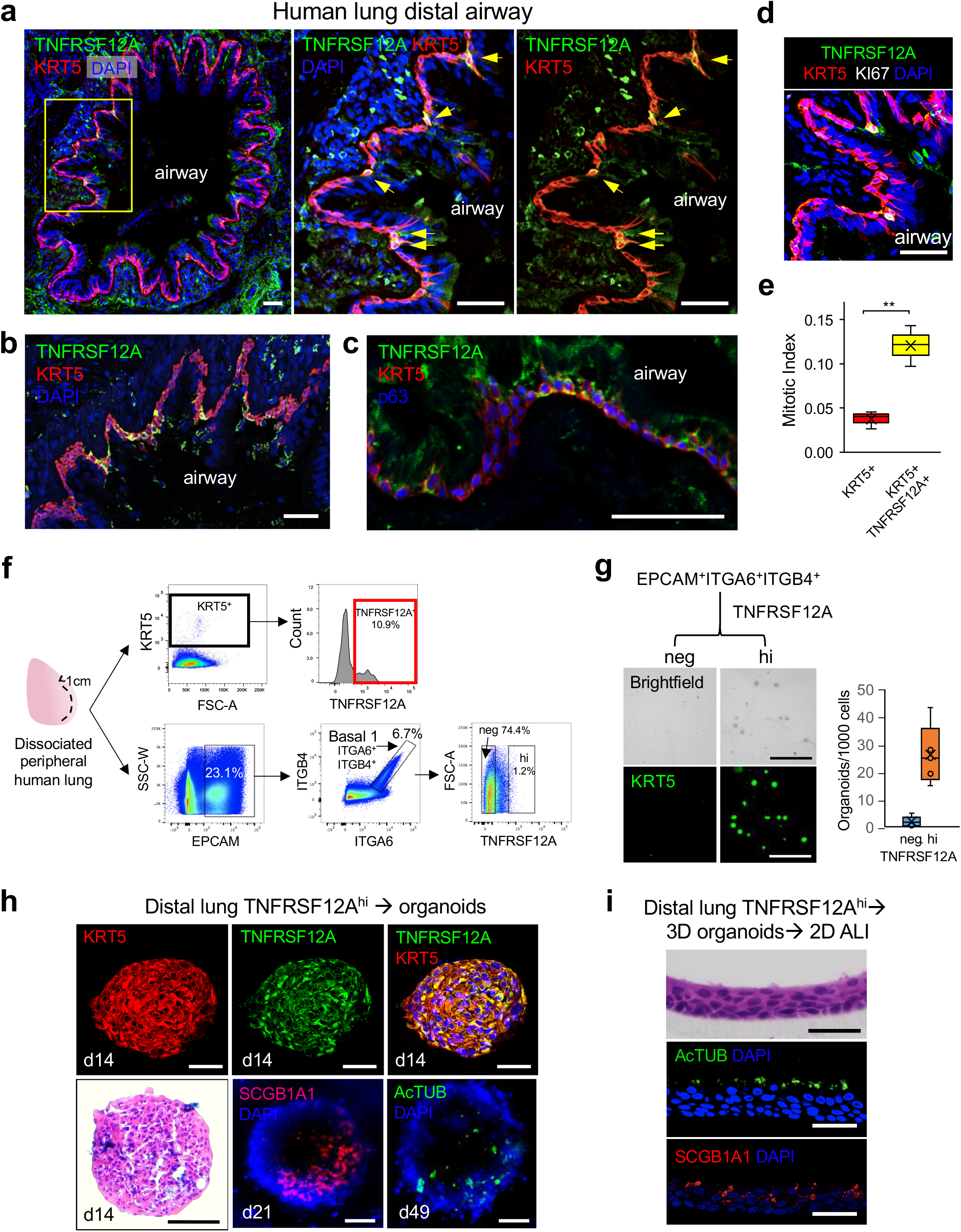
*In vivo* localization, prospective isolation, and clonogenic activity of the proliferative TNFRSF12A^hi^ subset of Basal 1 from intact human lung. **a,** Left, Representative multicolor immunofluorescence of freshly fixed human distal lung with overlap of monoclonal anti-KRT5 (red) and polyclonal anti-TNFRSF12A (green) in small airways, DAPI = blue. Middle, enlargement of the yellow boxed area from the left panel. Right, representation of the middle panel without DAPI channel, scale bar = 100 μm. **b,** Immunofluorescence of KRT5 (red) and TNFRSF12A (green) freshly fixed human distal lung small airways from an additional individual, scale bar = 100 μm. **c,** Representative immunofluorescence of TNFRSF12A (green), KRT5 (red), and p63 (blue) demonstrating TNFRSF12A overlap in a subset of KRT5^+^p63^+^ cells. **d,** Proliferation of TNFRSF12A^+^ airway basal cells in distal lung histologic sections. IF for KRT5 (red), TNFRSF12A (green) and KI67 (white) with DAPI (blue). scale bar = 100 μm. **e,** Mitotic index calculation of TNFRSF12A^+^ KRT5^+^ basal cells as in (b) from 3 biological replicates, error bars = SEM, * = p < 0.05 **f,** Schema outlining FACS analysis from freshly fixed human distal lung with anti-KRT5 (intracellular) and monoclonal anti-TNFRSF12A (cell surface) (top), or sequential FACS schema on viable cells freshly dissociated human distal lung to isolate EPCAM^+^ITGA6^+^ITGB4^+^ cells followed by fractionation into TNFRSF12A^hi^ or TNFRSF12A^neg^ subsets. All FACS populations were pre-gated on live singlets. **g,** Representative whole mount KRT5 staining and quantitation of clonogenic organoid formation from FACS-isolated TNFRSF12A^lo^ and TNFRSF12A^hi^ EPCAM^+^ITGA6^+^ITGB4^+^ cells as in (f) after 14 days culture, scale bar = 500 μm. Quantitation represents organoid growth per 1000 FACS isolated cells; data are from cultures from five unique individuals each with a mean of 3 technical replicates, * = p < 0.05. **h,** H&E and immunostaining of organoids generated from the TNFRSF12A^hi^ fraction of freshly dissociated EPCAM^+^ITGA6^+^ITGB4^+^ human distal lung cells. Whole mount staining for SCGB1A1 and acetylated tubulin (AcTUB) at the indicated culture time points. Scale bar = 50 μm. **i,** H&E and immunostaining of SCGB1A1 or acetylated tubulin (AcTUB) in 2D air liquid interface cultures initiated from basal cell organoids cultured from the TNFRSF12A^hi^ fraction of EPCAM^+^ITGA6^+^ITGB4^+^ freshly dissociated human distal lung cells. Scale bar = 50 μm.

TNFRSF12A^hi^ Basal 1 cells could be prospectively isolated and cultured directly from freshly dissociated human lungs for culture without an organoid intermediate. EPCAM^+^ITGA6^+^ITGB4^+^TNFRSF12A^hi^ (i.e. TNFRSF12A^hi^ Basal 1) and EPCAM^+^ITGA6^+^ITGB4^+^TNFRSF12A^neg^ cells (i.e. TNFRSF12^neg^ Basal 1) were isolated from dissociated human lungs **(Fig. 4f, bottom)** and cultured clonogenically **(Fig. 4g-i).** Compared to TNFRSF12A^neg^ populations, TNFRSF12A^hi^ cells purified directly from distal lung exhibited a robust 15-fold increase in KRT5^+^ organoid formation **(Fig. 4g)**, indicating substantial enrichment of organoid-generating capacity and recapitulating the enhanced clonogenic growth of organoid-derived TNFRSF12A^hi^ cells **(Fig. 3f-g, Extended Data 8a-d)**. Organoids grown from distal lung TNFRSF12A^hi^ cells also exhibited a characteristic basal histology and spontaneously differentiated to SCGB1A1^+^ club and ActTUB^+^ ciliated cells **(Fig. 4h)**. Further, distal lung TNFRSF12A^hi^-derived basal organoids adopted a typical stratified epithelial histology with apical ciliary and club cell differentiation upon replating as 2D air-liquid interface monolayers **(Fig. 4i)**.

### SARS-CoV-2 and influenza H1N1 infection of distal lung organoids

We next established the utility of distal lung organoids for human infectious disease modeling. Influenza virus strain H1N1 broadly injures both airway and alveolar epithelium^45^. Both basal and AT2 organoids from human distal lung cultures were avidly infected by a recombinant influenza H1N1 PR8 strain expressing GFP upon viral replication^46^ **(Fig. 5a, Supplementary Data 5)** and viral genomic RNA accumulated in mixed organoid culture supernatants over a 96 hr time course **(Fig. 5b),** similar to previous reports with human lung airway organoids^21^. Organoids also stained with lectins binding a2-3 and a2-6 sialic acid residues (**Extended Data Fig. 9a,b**), indicating functional influenza receptors as in intact human lung^47^. Pretreatment of lung organoids with zanamivir, which selectively targets influenza release from infected cells, did not inhibit H1N1 viral infection or replication, as shown by the lack of GFP attenuation. In contrast, the nucleoside analog 2’fluoro 2’deoxycytidine (FdC), which impairs replication across many virus families, was efficacious with EC_50_ consistent with previous reports^48^ **(Extended Data Fig. 9c)**. Screening of diverse antiviral compound classes in H1N1-infected organoids in a 48-well format assay, quantified by GFP, revealed differential effectiveness **(Extended Data Fig. 9d)**, suggesting utility for scalable therapeutics discovery.

**Figure 5.**
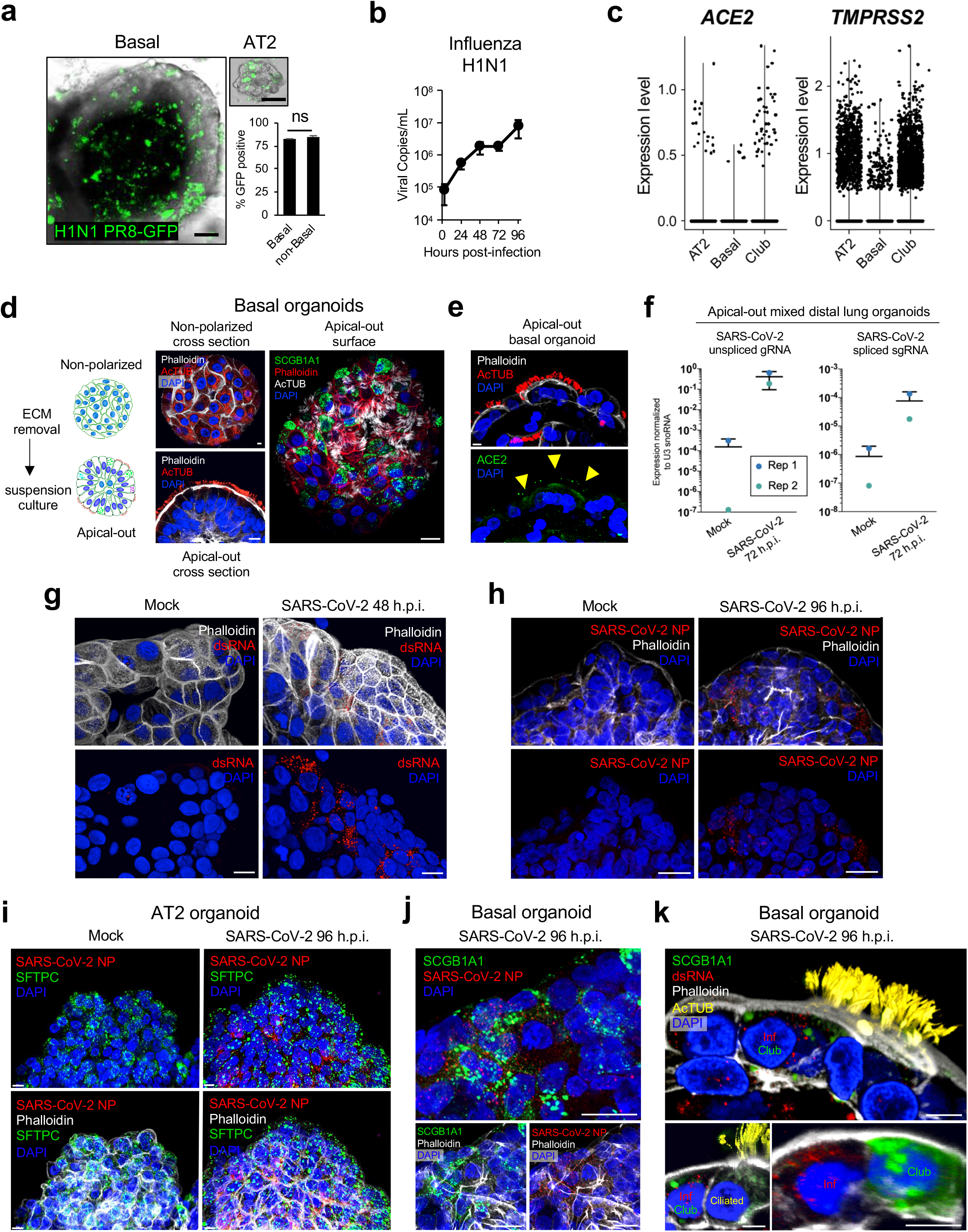
SARS-CoV-2 and influenza H1N1 infection of distal lung organoids. **a-b,** Mixed distal lung organoid modeling of H1N1 influenza infection. **a,** Merged transmission and GFP confocal images of purified basal (left) and purified AT2 organoids (right) 12 hours after infection with PR8-GFP H1N1 influenza, quantified by FACS for % GFP^+^ cells. Scale bars = 50 μm. High resolution images are provided in Supplementary Data. **b,** Viral genome quantitation over time of mixed distal lung organoid culture supernatants subjected to initial infection of wild-type H1N1 at an estimated multiplicity of infection (MOI) of 0.01, qRT-PCR, 3 biological replicates each with two technical duplicates, error bars = SEM, * = p <.05. **c,** scRNA-seq plots of *ACE2* and *TMPRSS2* gene expression in mixed distal lung organoids from Fig. 1q-r. **d,** Depiction of formation of apical-out lung organoids. Left: Diagram and representative confocal microscopy showing reorganization of microfilaments (phalloidin) and acetylated microtubules (AcTUB) upon ECM removal. Scale bar = 10 μm. Right: Confocal microscopy depicting differentiated cells (AcTUB^+^ ciliated cells and SCGB1A1^+^ club cells) exposed on the apical surface upon ECM removal. Scale bar = 20 μm. **e,** Confocal microscopy showing apical ACE2 immunofluorescence (yellow arrows) on apical-out basal organoids. Scale bar = 10 μm. **f,** qPCR of SARS-CoV-2 unspliced genomic RNA (left) and spliced subgenomic RNA (right) from infected apical-out distal lung organoids at 72 hours post-infection. n = 2 biological replicates. **g,** Confocal microscopy of double-stranded RNA (dsRNA) immunofluorescence on apical-out human distal lung organoids infected with SARS-CoV-2, mock vs. 48 hours post infection. Scale bar = 20 μm. **h,** Confocal microscopy demonstrating SARS-CoV-2 nucleocapsid protein (NP) immunofluorescence on apical-out human distal lung organoids infected with SARS-CoV-2, mock vs. 96 hours post infection. Scale bar = 20 μm. **i,** Confocal immunofluorescence analysis of SARS-CoV-2 infection of apical-out AT2 organoids with the indicated antibodies at 96 hours post-infection. Scale bar = 10 μm. **j,** Colocalization of SARS-CoV-2 NP and SCGB1A1 immunofluorescence 96 hours post infection of apical-out distal lung organoids. Scale bar = 20 μm. **k,** Cell type specificity of SARS-CoV-2 infection in apical-out distal lung organoids. Immunofluorescence was performed with the indicated antibodies. Top and bottom left: infected club cell adjacent to uninfected ciliated cells. Bottom right: infected cell adjacent to uninfected club cell. Inf = SARS-CoV-2 infected cell. Scale bar = 10 μm. In **e-k**, organoids were everted prior to infection for 6-10 days (basal) and 3 days (AT2).

The COVID-19 pandemic is particularly significant for distal lung disease, where involvement of alveoli and terminal bronchioles elicits life-threatening pneumonia and respiratory failure^1–3^. SARS-CoV-2 infects intestinal organoids^49–51^, and 2D ALI monolayer cultures from upper airway, trachea and alveoli^50,52^ but efficient infection of distal lung tissue has not been demonstrated. In mixed distal lung organoids, scRNA-seq **(Fig. 1q,r)** revealed mRNA encoding the SARS-CoV-2 receptor *ACE2* and the processing protease *TMPRSS2* predominantly in club and AT2 cells **(Fig. 5c),** consistent with ACE2 expression in KRT5^-^ differentiated lumenal cells **(Fig. 2j)**. The present long-term 3D basal and alveolar organoids are typically oriented with the basolateral surface oriented outwards, i.e. facing the extracellular matrix substratum, which could hinder SARS-CoV-2 infection of the apical ACE2-expressing luminal surface. We previously described rapid inversion of apical-basolateral polarity of gastrointestinal organoids by removal from the extracellular matrix gel and growth in suspension, robustly generating organoids with their apical surfaces oriented outward (apical-out), and thus greatly facilitating host-pathogen interactions on the lumenal surface^53^. We adapted this method to distal lung organoids where suspension culture rapidly induced apical-out polarization and differentiation. Within 48 h non-polarized KRT5^+^ organoids reorganized into apical-out epithelial spheroids with microvilli, apical junctions, and some motile cilia facing the organoid exterior. Within 5 days, differentiation of outwardly oriented ciliated cells continued to accelerate and was progressive over weeks **(Fig. 5d; Extended Data Fig. 10a-c; Supplemental Movie 3)**. Additionally, everted organoids displayed outwardly facing club cells with apical secretory granules **(Fig. 5d; Extended Data Fig. 10d**); delayed appearance of AT1 cells was also observed in alveolar organoids **(Extended Data Fig. 10e-g).** In apical-out organoids in suspension culture, ACE2 was detected on the apical membrane of cells on the external organoid surface **(Fig. 5e)** versus restricted to the internal differentiated apical lumen in basal organoids maintained in ECM **(Fig. 2j)**.

Notably, SARS-CoV-2 readily infected apical-out mixed distal lung organoids. SARS-CoV-2 genomic RNA was detected at 72 h post-infection by qPCR at levels similar to the abundantly expressed ubiquitous U3 snoRNA (Fig. 5f, left). Infection was further confirmed by the presence of replication-specific SARS-CoV-2 subgenomic RNA (sgRNA) (Fig. 5f, right) and by production of infectious virions capable of plaque formation on VeroE6 cells: 35 PFU/ml from cellular lysates and 65 PFU/ml from cellular supernatants. Immunofluorescence visualization of SARS-CoV-2-infected basal organoids revealed the sequential appearance of double-stranded RNA (dsRNA) by 48h, reflecting viral genome replication (Fig. 5g), and of SARS-CoV-2 nucleocapsid protein (NP) by 96h (Fig. 5h). We then examined which organoid cell types were infected by SARS-CoV-2 *in vitro*. Approximately 10% of AT2 organoids exhibited evidence of infection with prominent localization of SARS-CoV-2 NP within SFTPC-expressing cells (Fig. 5i). Similarly, SARS-CoV-2 infected ~10% of basal organoids. Since basal organoids contain multiple cell types, immunostaining of dsRNA or SARS-CoV2-2 NP was overlaid with either KRT5, SCGB1A1, or AcTUB to determine infected cell identity. In 2,621 total cells representing cultures from 4 individuals (Supplementary Data 7), SARS-CoV-2 infection was not detected in KRT5^+^ basal or AcTUB^+^ ciliated cells (odds ratio 0, p-value < 0.05), in contrast to prior studies in 2D ALI culture where SARS-CoV-2 infected upper airway ciliated cells^50,52^. However, SARS-CoV-2 NP and dsRNA immunofluorescence signals were primarily present in SCGB1A1^+^ club cells (Fig. 5j-k) which were strongly associated with and accounted for 79% of NP/dsRNA-positive cells (odds ratio 19.33, p < 0.0001); 21% of infected cells lacked SCGB1A1 (Fig. 5j-k, Supplementary Data 7). Overall, these studies indicated direct SARS-CoV-2 infection of AT2 cells, and implicated club cells as a novel target population.

## DISCUSSION

The slow cellular turnover of adult lung epithelium has hampered identification of regional stem cell populations. Lack of long-term culture systems and substantial differences between mouse and human lung have particularly impaired validation of human distal lung stem cells. Here, we described a robust, feeder-free, chemically defined method for long-term human distal lung airway and alveolar clonogenic organoid growth, which was applied to progenitor discovery and infectious disease modeling.

Basal cells perform crucial stem cell functions in lung and other tissues. Although lung molecular^40,54–57^ and histochemical^56,58^ basal cell subsets have been reported and differential progenitor activity has been observed in mouse^40,54^, proximal human airway studies suggest uniform basal cell proliferative activity *in vivo*^59^. We identified two interrelated molecular subtypes of KRT5^+^ human distal lung basal cells, Basal 1 and Basal 2; although we cannot exclude squamatization in the latter, proliferation selectively localized to the former. Notably, a TNFRSF12A^hi^ fraction within the Basal 1 subset, by both mRNA and protein criteria, possessed enriched clonogenic progenitor activity, establishing functional precedent for a proliferation-enriched basal cell subtype. TNFRSF12A^+^ basal cells often but not exclusively localized to bases and tips of airway furrows, possibly representing a distinct airway progenitor niche, as proposed for goblet cells^44^. Conceivably, TNFRSF12A or other markers could distinguish analogous human basal cell progenitor subsets in other tissues.

The differentiation of iPSC to lung epithelial lineages, while efficacious, can be constrained by efficiency, feeder dependence, and fetal gene expression^26–29^, necessitating methods to expand resident adult lung stem cells. Prior human lung basal cell cultures (nasal cavity, trachea, proximal bronchi) achieved in vitro proliferation and clonal expansion^16,60^, ciliated and mucous differentiation^16,41,60^ and reconstitution of denuded rat tracheal epithelium^58^ but have been limited by short term culture, feeder dependence, and restriction to upper airway^17,18^. Further, intrinsic differences in basal cells and their differentiated progeny along the lung proximal-distal axis may limit generalizability of upper airway studies to distal airways and alveoli^14^. Our distal airway basal organoids represent amongst the most significant clonal expansion of basal cells from any region of the human lung^14,23,24,60^.

Culture of human adult AT2 cells has been characteristically short-lived and feeder-dependent^6,18,24,25^. The present clonogenic, long-term, feeder-free and chemically defined human alveolar organoid cultures uniformly expressed the canonical AT2 genes *SFPTA1/B/C*, possessed characteristic lamellar bodies, functionally required autocrine WNT signaling^25,34^, and transdifferentiated to AT1 cells upon culture on glass or in suspension. Variable AT2 subpopulations were marked by *LYZ* or *MUC5B* **(Extended Data Fig. 5-6)**, previously identified in mouse and implicated in Idiopathic Pulmonary Fibrosis, respectively^61–63^. Murine lung progenitors with airway and alveolar epithelial differentiation capacity include LNEPs^9,64^, Distal Airway Stem Cells (DASCs)^10^ and Bronchioalveolar Stem Cells (BASCs)^9,10,12,65^. We did not observe clonal organoids simultaneously containing both mature airway and alveolar cell types but cannot exclude such bipotential progenitors upon different culture conditions.

We also describe a facile model for SARS-CoV-2 infection of distal lung, as relevant for COVID-19-associated pneumonia and ARDS^1–3^. Intestinal organoids recapitulate SARS-CoV-2 infection^49–51^, 2D ALI monolayer cultures from upper airway/trachea organoids are infected by SARS-CoV-2^50^ and a comprehensive study of 2D ALI cultures over the entire proximal-distal airway axis revealed a descending susceptibility gradient^52^. However, these studies neither employed 3D lung organoid systems nor demonstrated efficient infection of distal lung tissue. Previously, intestinal organoids were mechanically sheared to allow apical SARS-CoV-2 infection^50^. In contrast, we describe robust SARS-CoV-2 infection by everting distal lung organoids in suspension, rapidly creating physiologically relevant “apical-out” cultures that facilitate direct infection of the ACE2-expressing exterior apical surface that recapitulates *in vivo* physiology. Active SARS-CoV-2 infection of basal and AT2 organoids was evidenced by detection of spliced subgenomic viral RNA, dsRNA replication intermediates, viral nucleocapsid protein and infectious virus production. In addition to AT2 organoid infection, our results implicate SCGB1A1^+^ club cells as a novel SARS-CoV-2 distal lung target whose infection could compromise protective lung protective glycosaminoglycans, thus facilitating a vicious COVID-19 infection cycle. Although we did not observe ciliated cell infection, SCGB1A1-negative populations were also infected and are under further investigation; for example, bronchial transient secretory cells express *ACE2* and *TMPRSS2*^66^. Since SARS-CoV-2 infects cultured upper airway ciliated cells *in vitro*^50,52^, cognate distal lung ciliated cells could be less susceptible to SARS-CoV-2 or their infection facilitated by alternative culture conditions. These studies extend our prior pathogen investigations in apical-out GI organoids^53^, and the strong induction of functional ciliogenesis in apical-out suspension conditions may allow improved expansion of ciliated cells versus current 2D ALI monolayer protocols.

Overall, single cell analysis of organoid cultures, as exemplified here, may represent a general strategy for human stem cell investigation in slowly proliferating tissues. The culture of progenitors for all adult distal lung epithelial lineages, including alveoli, should substantially enable human pulmonary disease modeling including neoplastic and interstitial lung diseases ^67,68^ and allow tissue engineering and precision medicine applications. Finally, this organoid system should facilitate general mechanistic and therapeutic investigations of pulmonary pathogens, including the SARS-CoV-2 distal lung infection associated with fulminant respiratory failure.

## ACKNOWLEDGEMENTS

We are grateful to members of the Kuo and Desai labs for helpful discussions. We also thank the Stanford Tissue Bank, Joseph Shrager, Mark Berry and Winston Trope for tissue acquisition, the Stanford Stem Cell FACS Facility, Pauline Chu, Daniel Mendoza and Francisco de la Vega for technical expertise and James Zengel for SARS-CoV-2 primer design. SARS-Related Coronavirus 2, Isolate USA-WA1/2020, NR-52281 was deposited by the Centers for Disease Control and Prevention and obtained through BEI Resources, NIAID, NIH. A.A.S. was supported by the A.P. Giannini, ECOG-ACRIN Paul Carbone and Stanford Cancer Institute postdoctoral fellowships and NIH grant K08DE027730, and S.S.C. by the Stanford Medical Scientist Training Program. A.R. was supported by NIH grant T32 AI007502-23. J.Z. was supported by a Stanford Graduate Fellowship. S.M.D. by a CIRM Bridges Research Fellowship, R.A.F. by the Damon Runyon Cancer Research Foundation (DRG-2286-17), V.V.U. by the Netherlands Organization for Scientific Research, Rubicon grant (452181214), C.S. and J.Z. by NSF DMS 1712800 and the Stanford Discovery Innovation Fund, K.C.G. and M.M.D. by HHMI, and Burroughs Wellcome Fund Investigators in the Pathogenesis of Infectious Disease Grant 1016687 to C.A.B. T.J.D is the Woods Family Faculty Scholar and C.A.B the Tashia and John Morgridge Faculty Scholar in Pediatric Translational Medicine of the Stanford Maternal & Child Health Research Institute; C.A.B. is an Investigator of the Chan Zuckerberg Biohub. This work was also supported by NIH grants 5R01HL14254902 to T.J.D., U19AI057229 to M.M.D., U19AI116484, U01DK085527, U01CA217851, U01CA176299 and U01DE025188 to C.J.K., and California Institutes for Regenerative Medicine grant DISC2-09637 to C.J.K. and T.J.D., and the Bill and Melinda Gates Foundation OPP1113682 to C.J.K., M.A.R, and C.A.B.

## AUTHOR CONTRIBUTIONS

A.A.S. and S.S.C. conceived, designed, and performed experiments, analyzed data, and wrote the manuscript. A.R. and C.E.E. designed and performed SARS-CoV-2 infections, J.Z., V.V.U. and C.S. designed and interpreted single cell RNA-Seq studies. R.A.F. performed qRT-PCR. M.M.-C., S.M.D., A.J.M.S, T.U, J.J. and A.B. performed organoid culture and analysis. L.E.W. and M.M.D. designed FACS panels. V.L. and K.C.G. contributed the DLL4 E1E2 mutant. B.A. and S.P. performed SPADE analysis. K.N. and J.S.G. designed and executed influenza studies. G.X.Y.Z., J.M.T., P.B., S.B.Z. and T.S.M. were involved in single-cell RNA-seq experiments. P.H. provided in situ hybridization protocols. R.S.B. designed SARS-CoV-2 studies. M.R.A., C.A.B., T.J.D. and C.J.K. conceived and designed experiments, analyzed data, and wrote the manuscript.

## AUTHOR INFORMATION

Reprints and permissions information are available upon requesttoauthors. The authors declare competing financial interests. Correspondence and requests for materials should be addressed to, or.

## METHODS

### Human tissue procurement and processing

All material used in this work was approved by the Stanford School of Medicine’s Institutional Review Board and performed under protocol #28908. Standard informed consent for research was obtained in writing prior to tissue procurement. Peripheral lung tissue within 1 cm of the visceral pleura was obtained from surgical discards from lobectomies. For patients with suspected lung cancer, cases with clinical T4 (American Joint Cancer Committee 6^th^ edition) disease (e.g. features such as bronchial invasion or parenchymal satellite nodule/ metastases) were not used. Normal tissue was harvested from the lung margin most anatomically distal to palpably well-defined lesions, or from uninvolved lobes in the case of pneumonectomies. Samples with tumors containing ill-defined margins were deferred. Tissue was either processed fresh or placed at 4°C overnight and processed the following morning.

### Organoid culture

To isolate distal airway cells, lung parenchyma 1 cm from the visceral pleura was mechanically dissociated with Castro scissors, washed and incubated with 5 Units/ml porcine elastase (Worthington), 100 Kunitz Units/ml DNase I (Worthington), and Normocin (InvivoGen) resuspended in two tissue volumes of lung organoid media, comprised of Advanced DMEM/F12 (Invitrogen) supplemented with 10 mM nicotinamide, n-acetyl cysteine, 1X B27 supplement minus vitamin A, recombinant human NOGGIN (100 ng/ml, R&D Systems), recombinant human EGF (50 ng/ml, R&D Systems), and TGF-beta inhibitor A83-01 (100nM, Tocris). The tissue was then agitated for one hour at 37C and the resultant cell suspension was filtered through 100 through 40 μm cell strainers and subjected to ammonium chloride red blood cell lysis. The cell pellet was then washed and resuspended in 10 volumes of reduced growth factor Basement Membrane Extract II (Trevigen). Cells in matrix were then plated in 24-well plates in 50 microliter droplets, and warm media was added after the droplets solidified for ten minutes at room temperature Media was changed every 3-4 days and organoids were passaged every 3-4 weeks by dissociation with TrypLE. Passaging was based on ECM durability/integrity and estimated organoid confluency, judged by estimated organoid volume to volume of the ECM droplet. To rule out contamination by malignant cells, long-term cultures were systematically evaluated for the presence of dysplasia or carcinoma by a board-certified pathologist. In addition, five long-term organoid cultures (2-6 months) underwent targeted Next Generation Sequencing to detect the presence of pathogenic variants (see below). Full details are provided in **Supplementary Methods.**

### Tandem MACS stromal depletion and EPCAM purification of distal lung cells

Distal lung was dissociated as above, and all incubation steps were carried out on ice. 10^7^ cells were incubated with Fc Block (Biolegend 422301) and diluted 1:100 in FACS buffer (2 mM EDTA and 0.2% fetal calf serum in 1X PBS pH 7.4), for ten minutes followed by APC conjugated anti-CD45 antibodies at 1 μg/ml in FACS buffer for 30 minutes, washed, and subjected to two rounds of depletion with magnetic beads according to manufacturer’s protocol (Miltenyi: anti-human fibroblast 130-050-601, anti-CD31 130-091-935, anti-APC 130-090-855, LS column 130-042-401). Unlabeled cells were then centrifuged at 300 x *g* and labeled with a cocktail of 1 μg/ml of PerCP-Cy5.5 anti-EPCAM antibody and Zombie Aqua viability stain (Biolegend 423101) diluted 1:400 from stock concentration in FACS buffer.

### Organoid Cryopreservation and Recovery

For cryopreservation and recovery, ECM droplets were dissociated by pipetting in 3 volumes of PBS with 5 mM EDTA and then incubated on ice for one hour. Cells were pelleted at 300 x *g* for 5 minutes and resuspended in freezing medium (fetal bovine serum (Gibco), 10% v/v DMSO), placed into cryovials and then into Mr. Frosty™ (Thermo Fisher) containers and stored in a −80C freezer overnight, followed by transfer to liquid nitrogen vapor phase for long term storage. Organoids were recovered by quick thaw in a 37C water bath followed by washing in organoid media and plating in ECM with organoid media plus 10 μM ROCK inhibitor Y-27632 (Tocris).

### Screening exogenous growth factors in organoid culture

Distal airway cells were isolated and plated as above with the following exceptions: ADMEM/F12 was used instead of organoid medium during elastase digestion of lung tissue, cells were serially diluted and filtered through a 40 micron cell strainer and counted with a hemocytometer. 1000 viable epithelial cells (by Trypan blue exclusion, size, and morphology) per μL ECM were plated per 5 μL Matrigel droplet per well. Base media consisted of organoid media lacking A83-01, EGF, NOGGIN, WNT3A or RSPO1. EGF (final 50 ng/ml, R&D), NOGGIN (final 100 ng/ml, R&D), WNT3A (final 100 ng/ml, R&D), RSPO1 (final 500 ng/ml, Peprotech) or the PORCUPINE inhibitor C59 (final 1 μM, Biogems) were added singly or in combination to base media. Images were obtained ten days after primary plating with an inverted light microscope at 5X magnification. Each condition was plated in quadruplicate and organoid formation was quantified using the analyze particle (threshold = 490^2^ pixels) plugin in ImageJ as previously described^65^.

### Single Cell RNA-seq of unfractionated organoid cultures

Lung organoid cultures from separate individuals were dissociated 4 weeks after primary plating and subjected to droplet based scRNA-seq with the 10x Genomics Gemcode Single Cell 3’ platform with a 5 nucleotide UMI according to manufacturer’s protocol. Cell capture, library preparation, and sequencing were performed as previously described^69^. Principle Component Analysis, t distributed Stochastic Neighborhood Embedding, unsupervised Graph based clustering, statistical testing for all scRNA-seq analyses are described in **Supplementary Methods** and **Supplementary Data 3**.

### Single cell RNA-seq of purified AT2 organoid cultures

LysoTracker^+^ AT2 cells from unfractionated organoids were purified by FACS and cultured for two months with one passage. These were dissociated and subjected to droplet-based scRNA-seq with the 10x Genomics Chromium Single Cell 3’ platform v2 according to the manufacturer’s protocol. The library was sequenced using paired-end sequencing (26bp Read 1 and 98bp Read 2) with a single sample index (8bp) on an Illumina NextSeq 500. Data preprocessing and Principle Component Analysis were carried out with CellRanger v1.2. Subsequent analysis is described in **Supplementary Methods** and **Supplementary Data 3**.

### Electron microscopy

Organoid cultures were fixed in ECM with 2.5% glutaraldehyde in 0.1 M cacodylate buffer (pH 7.4), dehydrated, embedded in epoxy resin and visualized with a JEOL (model JEM1400) transmission-electron microscope with a LaB6 emitter at 120 kV.

### Histology and immunocytochemistry

Organoids were fixed with 2% paraformaldehyde at 4 degrees Celsius overnight, paraffin embedded and sectioned (10-20 μm) as previously described^69^. Sections were deparaffinized and stained with H&E for histological analysis. Antibodies used for immunocytochemistry staining are listed in **Supplementary Methods** following standard staining protocol as previously described^70^ and images were acquired on a Leica-SP8 confocal microscope.

### RNA fluorescent in situ hybridization

RNA in situ hybridization was performed according to Nagendran et al.^71^ and probe sequences are provided in **Supplementary Methods**.

### Whole mount organoid confocal immunofluorescence microscopy

Intact, uninfected organoids were fixed in 2% paraformaldehyde in 100 mM phosphate buffer (pH 7.4) (4% paraformaldehyde for infected organoids) for one hour at room temperature, washed with PBS with 100 mM glycine, permeabilized 0.5% triton X-100 in PBS for one hour, then incubated in staining buffer (4% BSA, 0.05% Tween-20 in PBS pH7.4, 10% goat/donkey serum) for an additional hour, followed by incubation with primary antibody for 24 hours at room temperature in staining buffer. Whole mounts were then washed with PBS-T and incubated with fluorescent secondary antibodies, phalloidin and DAPI, for four hours at room temperature in staining buffer. Following additional washes, whole mounts were submerged in mounting media (VECTASHIELD, Vector Laboratories) and mounted on chambered coverslips for imaging in four channels using Zeiss LSM 700 or 900 confocal microscopes. 3D rendering of confocal image stacks was performed using Volocity Image Analysis software (Quorum Technologies Inc., Guelph, Ontario). For Figure 5j, requiring 5 colors, cilia were distinguished by staining with two fluorescent secondary antibodies and merging the colocalized voxels into a pseudocolored channel using Volocity software. Lectin staining (FITC-Sambuca Nigrin, Vector Labs FL-1301; Biotin-Maackia Amurensis, Vector Labs FL-1301) was carried out according to manufacturer’s protocol after fixation of organoids with 0.1% paraformaldehyde in PBS for 1 hour at room temperature followed by blocking with Avidin/Biotin (Vector Labs SP-2001). Biotin-Maackia Amurensis lectin was labeled with streptavidin-PE conjugate (Thermo Fisher SA10041) and after washing lectin staining was imaged in a Keyence BZ-X700.

### Next generation sequencing of organoid cultures

Ten organoid cultures were sequenced using a commercial targeted resequencing assay with end-to-end coverage of 131 cancer genes and companion software (TOMA COMPASS Tumor Mutational Profiling System, Foster City, CA) to determine the presence of oncogenic mutations in long-term organoid cultures. Libraries were sequenced on an Illumina NextSeq 500. Nonsynonymous variants are listed in **Supplementary Table 3**. Variant Call Files are provided in **Supplementary Data 6**.

### Density sedimentation of basal cells

Organoid cultures within 2-3 weeks of primary plating were dissociated with 1 U/ml neutral protease (Worthington, Cat LS02100) and 100 KU of DNase I in organoid media. Basal organoids were then collected by gravity sedimentation and the supernatant was either aspirated or collected for downstream use. Basal organoids were then further fractionated on a custom Ficoll-Paque gradient (4 vol Ficoll-Paque to 1 vol PBS) and centrifuged at 300 x *g* for 10 minutes at room temperature. The supernatant was aspirated and the organoid pellet was resuspended in 10 ml PBS in a 15 ml conical tube, collected by gravity sedimentation, and plated into ECM as described above.

### FACS isolation and culture of AT2 cells

Organoids were dissociated with TrypLE followed by neutralization with 10% volume fetal calf serum, subjected to DNase at 100 kU/ml, washed with organoid media and then incubated with 100 cell pellet volumes of organoid media with 10 nM LysoTracker Red DND-99 (Thermo Fisher L7528) at 37C for 30 minutes. Cells were then washed and resuspended in FACS buffer as described above, incubated with Fc block, followed by incubation on ice with labeling cocktail consisting of 1 μg/ml of PerCP-Cy5.5 anti-EPCAM antibody and Zombie Aqua viability stain (Biolegend 423101) diluted 1:400 from stock concentration in FACS buffer. EPCAM^hi^ and LysoTracker^hi^ cells were sorted into organoid media with 10 μM Y-27632 (Tocris 1254) and cultured in ECM and media with Y-27632 for 24 hours, followed by regular media. Full gating strategy is provided in **Supplementary Data 2**. All FACS antibodies were purchased from Biolegend.

### Color mixing studies with lentivirally transduced GFP and mCherry

FACS EPCAM^+^ stromal depleted organoids at d14 were infected with lentivirus at an estimated MOI of 0.9 according to Van Lidth de Jeude et al.^72^ with third generation lentiviral vectors (PGK-GFP T2A Puro, SBI cat# CD550A-1; mCherry modified from pLentiCRISPRv1 (Addgene #49545) to incorporate an EF-1a-mCherry P2A Puro cassette, a gift from Paul Rack). 96 hours after infection, organoids were treated with puromycin at a concentration of 600 ng/ml for 48 hours to select for transduced cells. Two weeks after selection, GFP expressing organoids and mCherry expressing organoids were dissociated to single cells and mixed in a 1:1 ratio and scored as monochromatic or mixed after 28 days of each passage. The same approach was employed for purified AT2 and basal cultures after respective purification strategies from an initial FACS EPCAM^+^ stromal depleted organoid starter culture.

### Flow cytometry analysis of resident basal cells from adult human lung

Adult human lung tissue was procured and dissociated as above but cells were labeled with Zombie Aqua live:dead stain as above, washed with FACS buffer, and then fixed in 2% PFA in PBS overnight at 4C. Cells were then stained using the whole mount procedure as described above with the omission of PBS glycine washing. Fixed and permeabilized cells were then incubated with 1:400 dilution of Alexa Fluor 647 conjugated mouse anti-human cytokeratin 5 antibody (Abcam) for 24 hours at 4C in permeabilization buffer. Cells were then washed with FACS buffer and labeled with PE conjugated mouse anti-human TNFRSF12A antibody (clone ITEM-4, Biolegend) for 30 minutes on ice, followed by washing and analysis on a BD Aria Fusion instrument. Full gating strategy and qPCR validation of ITEM-4 antibody is detailed in **Supplementary Data 2**.

### FACS isolation of TNFRSF12A^hi^ and TNFRSF12A^neg^ basal cells

Single cell suspensions from either fresh human distal lung or primary organoid culture at approximately 4 weeks of culture were dissociated as above, treated with Fc Block, and incubated in FACS buffer with Zombie Aqua 1:400, 1 μg/ml PerCP-Cy5.5 anti-human EpCAM (CD326), 1 ug/ml APC anti-human ITGA6 (CD49f), 2 ug/ml FITC anti-human ITGB4 (CD104), and 1 μg/ml PE anti-human TNFRSF12A (CD266). 30 minutes after labeling the cells were washed twice with FACS buffer and sorted for EPCAM^hi^, ITGA6/ITGB4^hi^, TNFRSF12A^hi^ and TNFRSF12A^neg^. Full gating strategy is provided in **Supplementary Data 2**. > 5000 cells were sorted into Eppendorf tubes with lung organoid medium and 10 μM ROCK inhibitor Y-27632. All FACS antibodies were purchased from Biolegend.

### Culture of TNFRSF12A^hi^ and TNFRSF12A^neg^ basal cells

Cells were seeded in ECM and submerged in lung organoid media with 10 μM ROCK inhibitor Y-27632. Seeding density for cells FACS isolated from organoid culture was 1000 cells per well at a density of 100 cells/μL of ECM. Seeding for cells FACS isolated from fresh human distal lung was 3000 cells per well at a density of 300 cells/μL of ECM. After 24 hours, the media was changed to remove ROCK inhibitor and additionally changed every 72 hours. Organoid formation was manually quantified 14 days post plating by two independent observers.

### NOTCH manipulation in TNFRSF12A^hi^ basal cells

TNFRSF12A^hi^ basal cells were isolated and cultured as above. 24 hours after plating, media was changed to lung organoid media with either vehicle (0.1% DMSO), 1 μg/ml JAG1 peptide (Anaspec), 500 nM soluble recombinant NOTCH receptor inhibitor Delta Like Ligand 4 mutant (DLL4E12), or 1 μM gamma secretase inhibitor DBZ (Tocris). After 14 days of culture, biochemical estimation of proliferation was carried out by resazurin reduction assay (Alamar Blue, Thermo Fisher) for 16 hours according to manufacturer’s protocol and resazurin reduction was measured via fluorescence readout on a Biotek Synergy H1 plate reader according to the manufacturer’s protocol. Reference blank consisted of Alamar Blue reagent incubated in parallel media without cells. Expression and purification of the NOTCH receptor inhibitor DLL4E12 was performed as previously described^39^.

### H1N1 organoid influenza assay

Unfractionated cultures containing AT2, basal, and club cell types at 2-3 weeks were infected in triplicate with PR8 strain of H1N1 modified to express GFP upon viral replication^46^ after 24 hours of pretreatment with antiviral compounds. ECM was dispersed by addition of 5 mM EDTA in PBS, followed by washing and inoculation with GFP-reporter virus at an estimated MOI of 1 in media containing either vehicle or antivirals. After 12 hours (one influenza infection cycle), intact organoid GFP expression was visualized by either fluorescence microscopy with a Keyence BZ-X700 automated microscope, or dissociated to single cell, fixed with 0.1% PFA in PBS followed by flow cytometry quantitation of GFP^+^ cells (Gating Strategy is provided in **Supplementary Data 2**). Antiviral dose response curves were generated using four-parameter nonlinear regression curve fitting with GraphPad Prism 7 (GraphPad Software, San Diego, CA). H1N1 tropism was assessed in a manner similar to above with the exception of Ficoll sedimented basal cell fraction versus non-basal fractions were dissociated to single cells, counted, and infected with an estimated MOI of 1 in organoid media for one hour at 37°C, followed by washing and reseeding into ECM, cultured for 16 hours, followed by dissociation and flow cytometry analysis as above.

### Quantifying H1N1 infection productivity

Productivity of pandemic H1N1 virus infection (A/California/07/2009) was determined by qPCR according to Zhou et al^21^ but with the following modifications. Organoids at 6 weeks of culture were removed from ECM with 1 U/ml neutral protease, washed with media, and reseeded 1:1 in 24 well plate wells 10% ECM and organoid media for 24 hours. Virus was added at an estimated MOI of 0.01 and incubated for 2 hours at 37°C. The supernatant was removed and wells were washed thrice with media and incubated with 1 ml of media per well. 250 μL aliquots of cell culture supernatant were harvested at 2, 24, 48, 72 and 96 hour time points with an equal volume of media replaced for each aliquot. Virus was quantified by qPCR according to Krafft et al^73^.

### Suspension culture to generate apical-out polarity in lung organoids

Lung organoids grown embedded in 50μl ECM-droplets were transferred to suspension culture as described in Co et al^53^ with some modifications. Briefly, ECM-embedded organoids were dislodged gently by pipetting using sterile LoBind tips (Eppendorf 22493008) and placed in 15 ml LoBind conical tubes (Eppendorf 30122216) containing ice-cold 5 mM EDTA in PBS. The ratio of EDTA solution to ECM and the time of solubilization is important to optimally release intact organoids from the matrix. 5 ml of EDTA solution are used per ECM-droplet (3 ECM droplets/15 ml conical) rotating for 1 h at 4°C on a rotating platform. Organoids were centrifuged at 200 x g for 3 min at 4°C and the supernatant was removed. The pellet was re-suspended in growth media in ultra-low attachment 6-well tissue culture plates (Corning Costar 3471). Suspended organoids were incubated at 37°C with 5% CO2 for different times (range 0-30 days) to characterize apical-out polarity, ciliogenesis, and differentiation, and to prepare apical-out organoids for infection experiments with SARS-CoV-2.

### SARS-CoV2 infection of human distal lung organoids

VeroE6 cells were obtained from ATCC and maintained in supplemented DMEM with 10% FBS. SARS-CoV-2 (USA-WA1/2020) was passaged in VeroE6 cells in DMEM with 2% FBS. Titers were determined by plaque assay on VeroE6 cells using Avicel (FMC Biopolymer) and crystal violet (Sigma), viral genome sequence was verified, and all infections were done with passage 3 virus. Organoids were counted and passaged into suspension media for 6-8 days and then resuspended in virus media or an equal volume of mock media, at a MOI of 1 relative to total organoid cells in the sample, and then incubated at 37°C under 5% CO2 for 2 hours. Organoids were then plated in suspension in EN media (apical-out organoids). At the indicated timepoints, organoids were washed with EN media and PBS and either resuspended in TRIzol LS (Thermo Fisher), freshly-made 4% PFA in PBS, or 250 μL EN media. Cells resuspended in EN media were lysed by freezing at −80. Culture supernatants were preserved in TRIzol LS or added directly to plaque assay monolayers. All SARS-CoV-2 work was performed in a class II biosafety cabinet under BSL3 conditions at Stanford University.

### qPCR analysis of SARS-CoV-2 RNA

RNA from SARS-CoV-2-infected organoids was extracted by adding 750 μl TRIzol (Thermo Fisher Scientific), incubating at 55 °C for 5 min and then adding 150 μl chloroform. After mixing each sample by vortexing for 7 s, the samples were incubated at 25 °C for 5 min and then centrifuged at 12,000 r.p.m. for 15 min at 4 °C. The aqueous layer was carefully removed from each sample, mixed with two volumes of 100% ethanol and purified using an RNA Clean & Concentrator-25 kit (Zymo Research) as per the manufacturer’s instructions. All RNA samples were DNase treated with the Turbo DNA-free kit (Thermo Fisher Scientific). The Brilliant II SYBR Green QRT-PCR 1-Step Master Mix (VWR) was used to convert RNA to cDNA and amplify specific RNA regions on the CFX96 Touch real-time PCR detection system (Bio-Rad). RT reaction was performed for 30 min at 50 °C, 10 min at 95 °C, followed by two-step qPCR with 95 °C for 10 seconds and 55 °C for 30 seconds, for a total of 40 cycles. Two primer sets were used, either to amplify non-spliced SARS-CoV-2 genomic RNA (gRNA) spanning nucleotide positions 14221-14306, or spliced SARS-CoV-2 sgRNA^52^. The primer sequences are listed in the Supplementary Methods.

### Quantitation of *SCGB1A1* and *SFTPC* mRNA expression in TNFRSF12A^hi^ basal cells

FACS-isolated basal cells cultured in the above conditions were fixed in 2% paraformaldehyde in PBS for one hour at room temperature, embedded in HistoGel (Thermo Fisher HG-4000-012), dehydrated and paraffin embedded en bloc. One hundred serial sections were obtained at 10 micron thickness, and immunocytochemistry and in situ hybridization were performed at each 100 micron level. Confocal images were acquired in a blinded manner and organoids were defined as a cluster of 3 or greater DAPI nuclei. Channels were acquired using identical parameters for DAPI, Alexa 488 (*SFTPC* RNA in situ hybridization), Texas Red (*SCGB1A1* RNA in situ hybridization), and Alexa 647 (KRT5 immunostaining). Z-stacks were collected and images were processed in ImageJ and maximum intensity Z projections were used to quantitate *SCGB1A1* and *SFTPC* RNA in situ in Cell Profiler^74^ using the RNA Proximity Ligase Assay counting pipeline (https://github.com/tischi/cellprofiler-practical-NeuBIAS-Lisbon-2017).

### TNFRSF12A immunostaining of intact distal lung

Optimal staining of human distal lung tissue was achieved from specimens fixed within 30 minutes of primary surgical resections in 4% paraformaldehyde in PBS. Specimens were incubated in fixative overnight at 4°C, transferred to 30% sucrose, and embedded into OCT. 10 μm thick frozen sections were cut, subjected to citrate based antigen retrieval (Vector labs) at 70°C for 30 minutes, followed by blocking for one hour with 10% goat serum in IF wash buffer as described above. Mouse anti-TNFRSF12A (clone ITEM-4, Biolegend) was utilized for **(Fig. 4f-i)** and polyclonal rabbit anti-TNFRSF12A (ThermoFisher PA5-20275) was used for **Fig. 4a-e, h)**.

### Live-imaging and confocal microscopy of immobilized apical-out lung organoids

Live organoids were held between two coverslips in a viewing chamber (Lab-Tek II two-chambered coverglass) and filmed using a Nikon TE2000E microscope using differential interference contrast (DIC) microscopy with a 63X objective. Samples were kept at 37°C with 5% CO2 during imaging. Digital videos were collected by a Hamamatsu high-resolution ORCA-285 digital camera and rendered using OpenLab 5.5.2 software (Improvision). After recording, samples were fixed and stained without removal from the chambers and transferred to the confocal microscope for immunofluorescence microscopy.

### Additional experimental details are in Supplementary Methods

