## Extended_Data for "Progenitor identification and SARS-CoV-2 infection in long-term human distal lung organoid cultures"

### Extended Data Figure 1

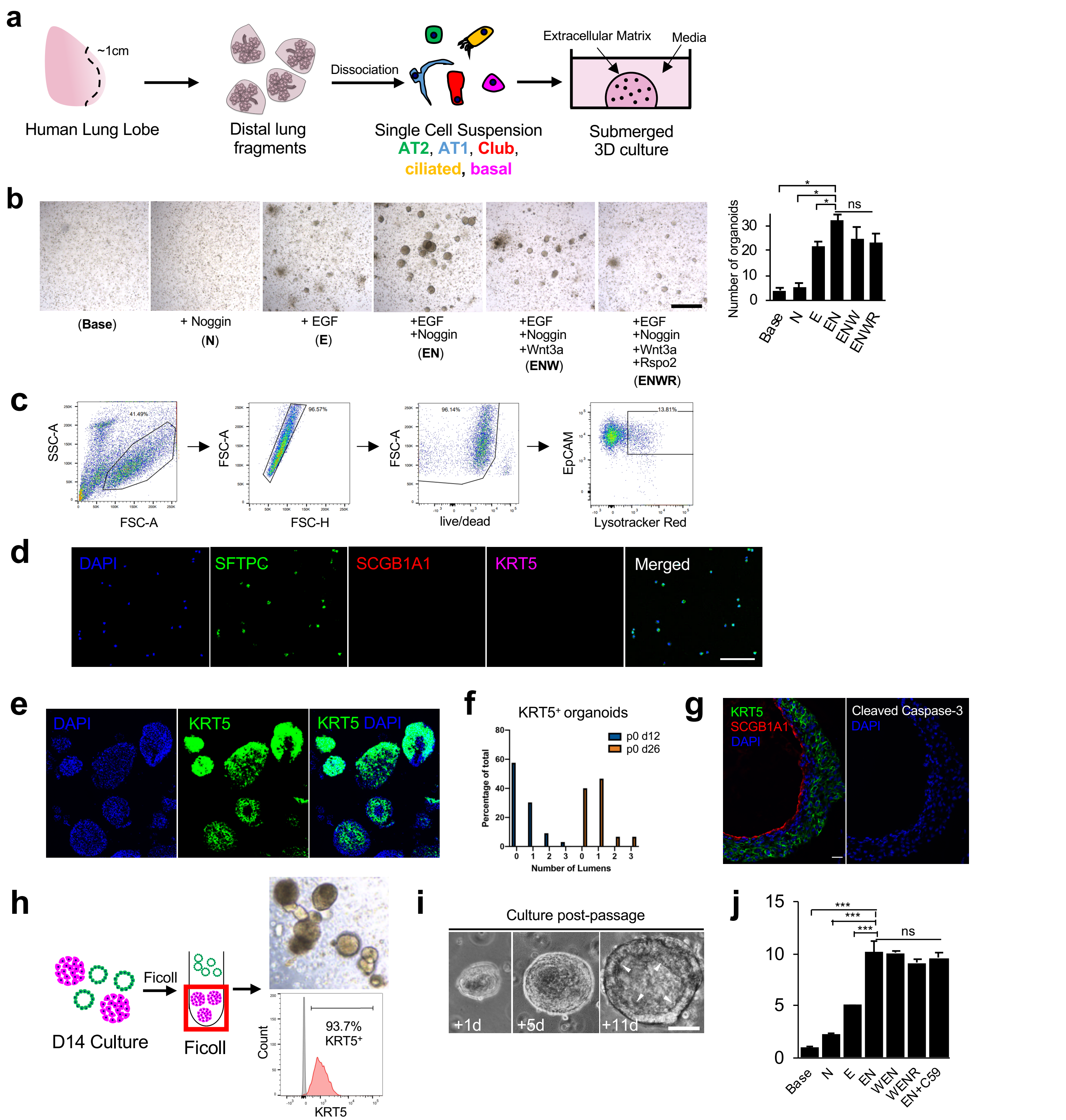


**Extended Data Figure 1.** Optimization of organoid culture of human distal lung. **a,** Schematic of culture initiation from human distal lung. **b,** Brightfield microscopy evaluation of required exogenous growth factors and automated organoid quantitation. **c-d,** Isolation of purified AT2 organoids. **c,** Representative FACS plots showing AT2 purification from unfractionated organoid cultures. **d,** Immunostaining of cytospin of sorted AT2 cells from (c) show high purity (100/100 cells SPC$^+$ SCGB1A1$^-$ KRT5$^-$); scale bar = 50 μM). **e-g,** Basal organoids in mixed culture progressively form internal lumens which is not associated with apoptosis. e, KRT5 IF, day 26 culture. **f.** Lumen quantitation, d12 versus d26 culture. **g,** Absence of apoptosis in d26 basal cell organoid internal lumen, cleaved caspase IF, from Fig. 2k, scale bar = 20 μm. **h-j,** Isolation of purified basal cell organoids via differential sedimentation in Ficoll. **h,** Schema and enrichment to > 90% KRT5$^+$ cells as measured by intracellular KRT5 FACS of sedimented basal organoid cells; scale bar = 100 μm. **i,** Serial time lapse microscopy of sedimented basal organoids reveals spontaneous cavitation within two weeks post passage or within four weeks of culture initiation; scale bar = 25 μm. **j,** Growth factor evaluation for basal organoids after d14 sedimentation, enzymatic dissociation and clonogenic culture. Growth was not affected by the PORCUPINE inhibitor C59 (1 μM). n=3 technical replicates, error bars = SEM, * = $p < 0.05$.

### Extended Data Figure 2

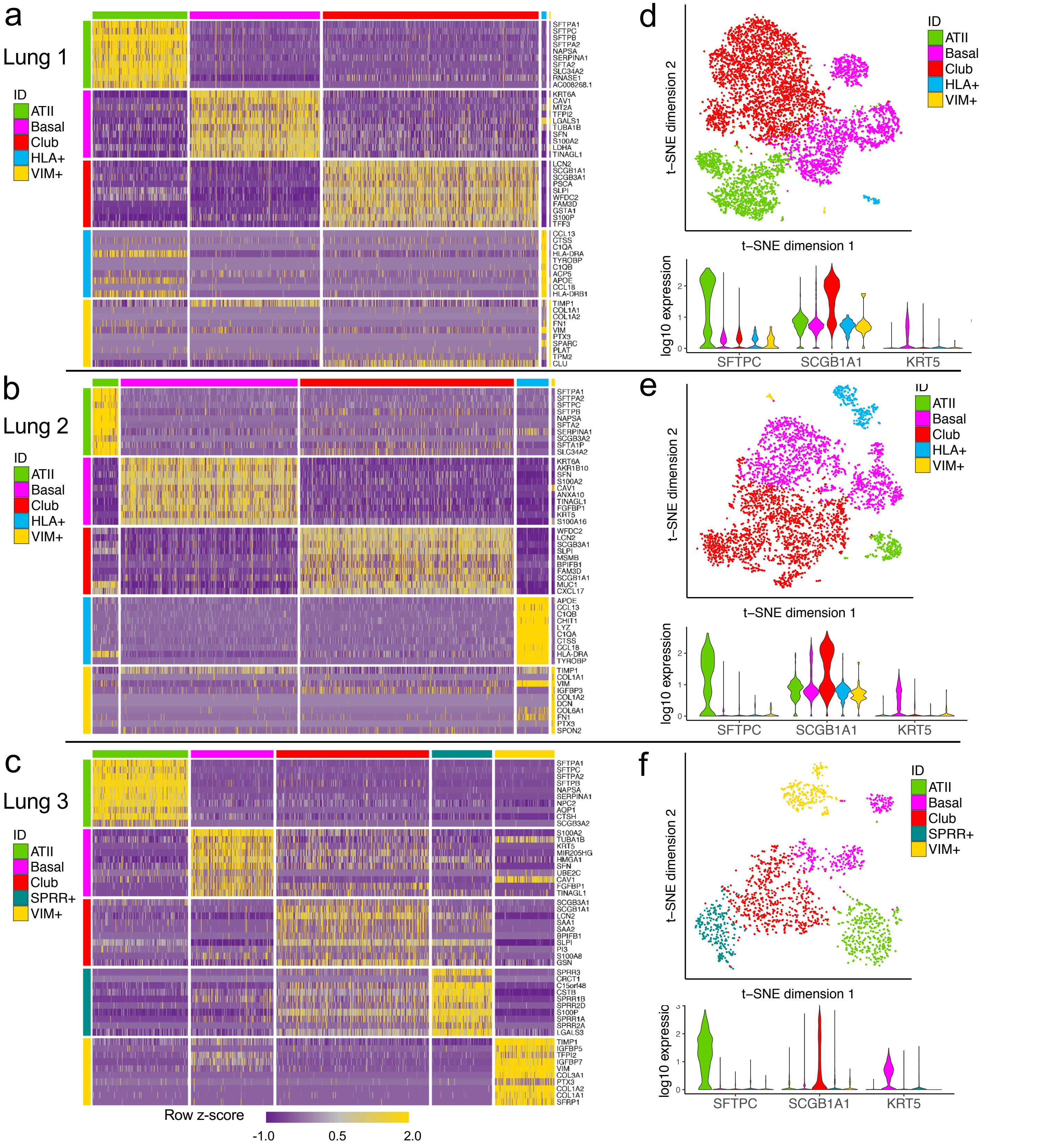


**Extended Data Figure 2.** scRNA-seq of human distal lung organoids reveals reproducible basal, club, and AT2 populations cultured from three individuals. **a-c,** unsupervised clustering of total cell populations demonstrates consistency in top differentially expressed genes corresponding to basal (*KRT5/6*), club (*SCGB1A1*), and AT2 (*SFTPC*) cells. The epithelial fraction from these cultures ranged from approximately 90-99% of all cells with the remainder being either fibroblasts (*VIM+*) or mononuclear cells (*HLA+*, likely alveolar macrophages). **d-f,** t-SNE visualization and violin plots for marker genes corresponding to each population. Note, a unique population enriched for SPRR genes, which have been described as a marker in squamous metaplasia, were exclusively found in the organoid culture of Lung 3, derived from an individual who was an active smoker.

### Extended Data Figure 3

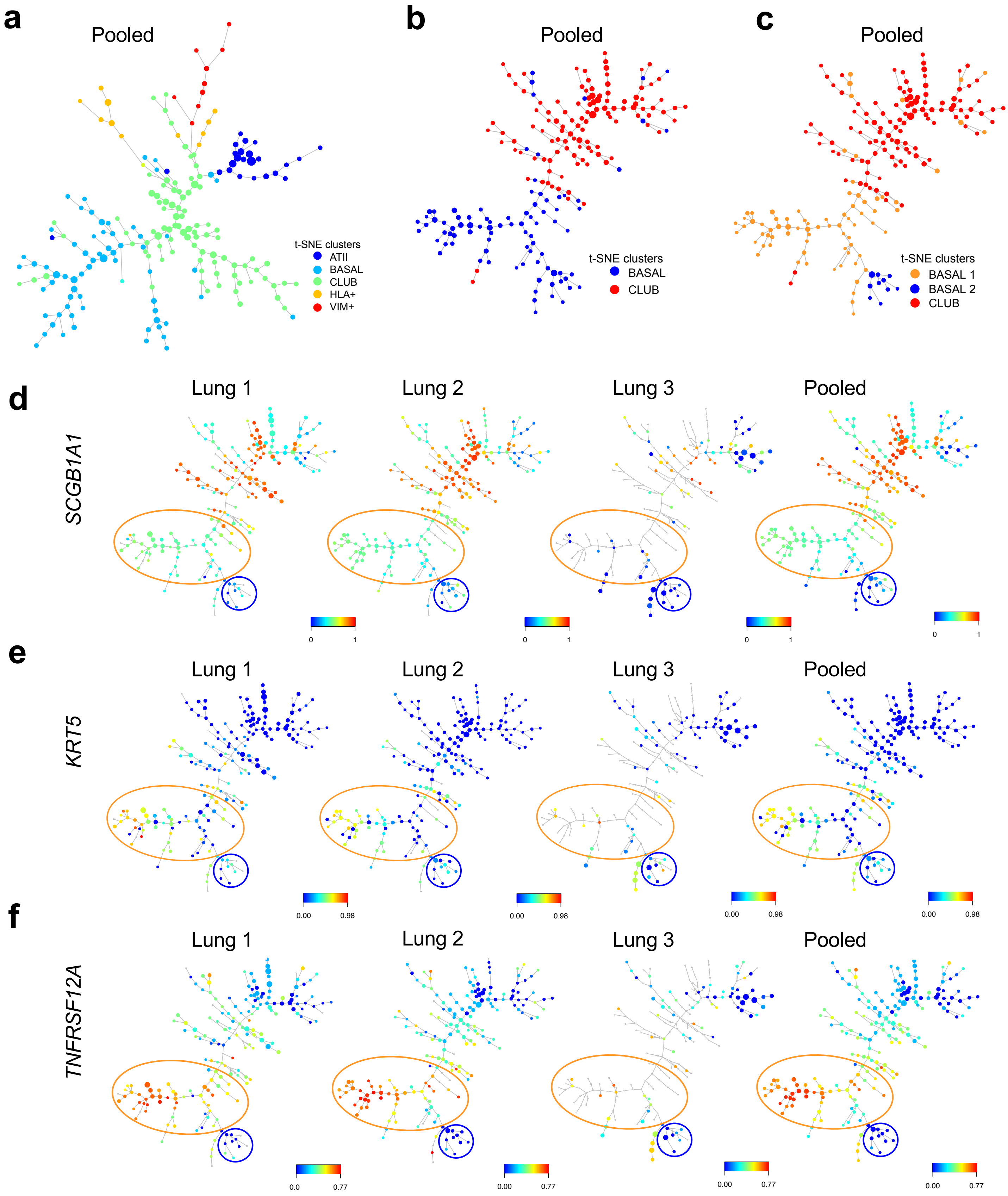


**Extended Data Figure 3.** Trajectory inference with SPADE. **a,** SPADE plot of cells where each point represents cell states that are more related on the same or adjacent branches of a minimum spanning tree. Note: AT2 cells exist on a branch distal to basal and club cells, suggesting no lineage hierarchy between AT2, basal, and Club cells. **b,** SPADE plots of pooled scRNA-seq samples after excluding AT2, *VIM*$^{+}$ and *HLA*$^{+}$ cells support lineage relationships between basal (blue) and Club (red) populations by Club cell branches emanating from basal cells. **c,** SPADE plots of Basal 1, Basal 2, and Club populations. **d,** gene expression of *SCGB1A1* shows higher expression in Club versus basal cell lineages as compared to **e,** KRT5. **f,** Median gene expression of *TNFRSF12A*, showing a high (orange outline) and a low (blue outline) within basal cell branches and inferring a potential lineage relationship.

### Extended Data Figure 4

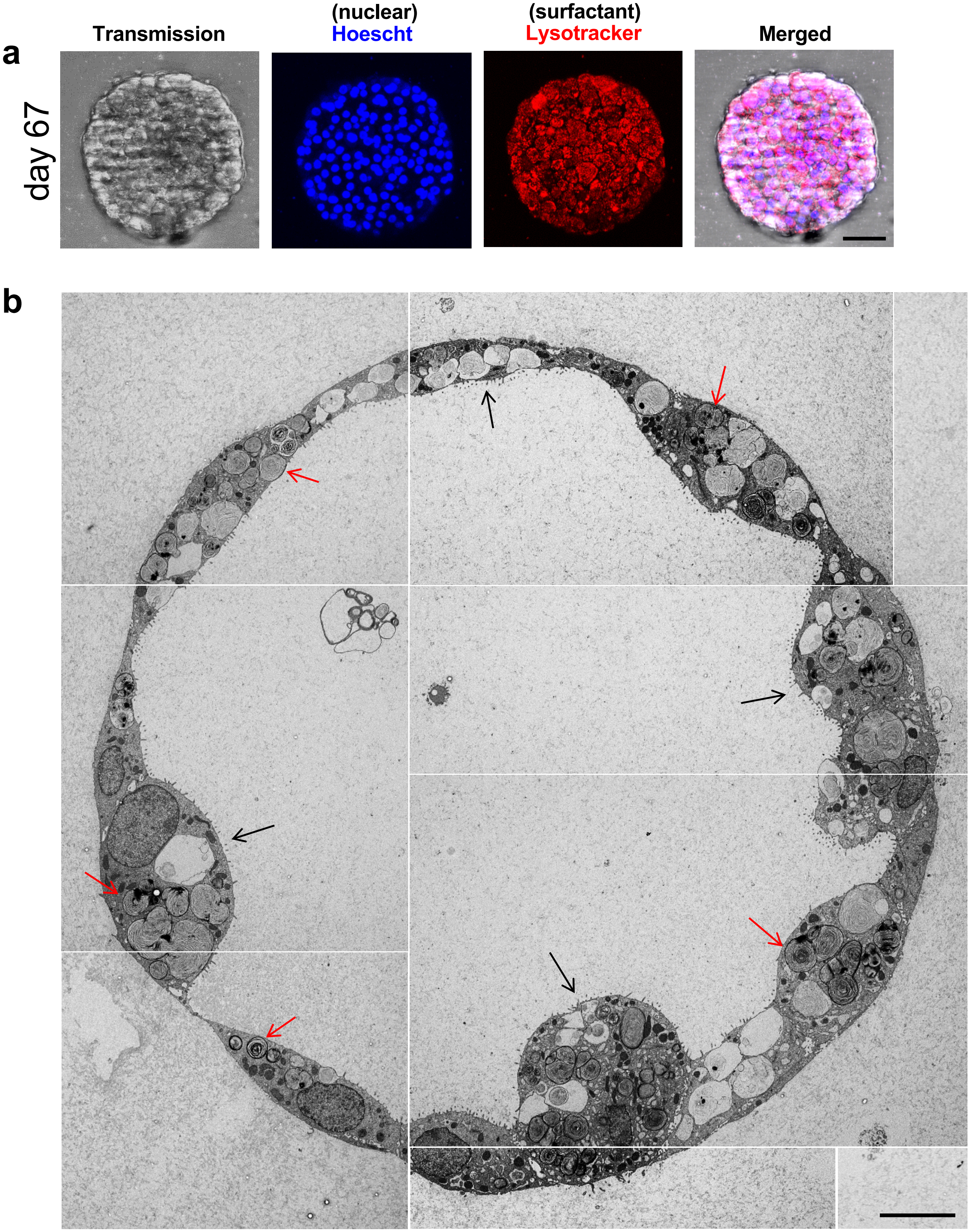


**Extended Data Figure 4.** Human AT2 organoid cells contain lamellar bodies. **a,** Confocal images of a live AT2 organoid at 67 days of culture labeled with Hoescht nuclear stain and LysoTracker Red DND-99. **b,** Transmission electron microscopy image of representative AT2 organoid at 28 days of culture. Note apical microvilli (black arrows) and lamellar bodies (red arrows); scale bar = 10 μm.

### Extended Data Figure 5

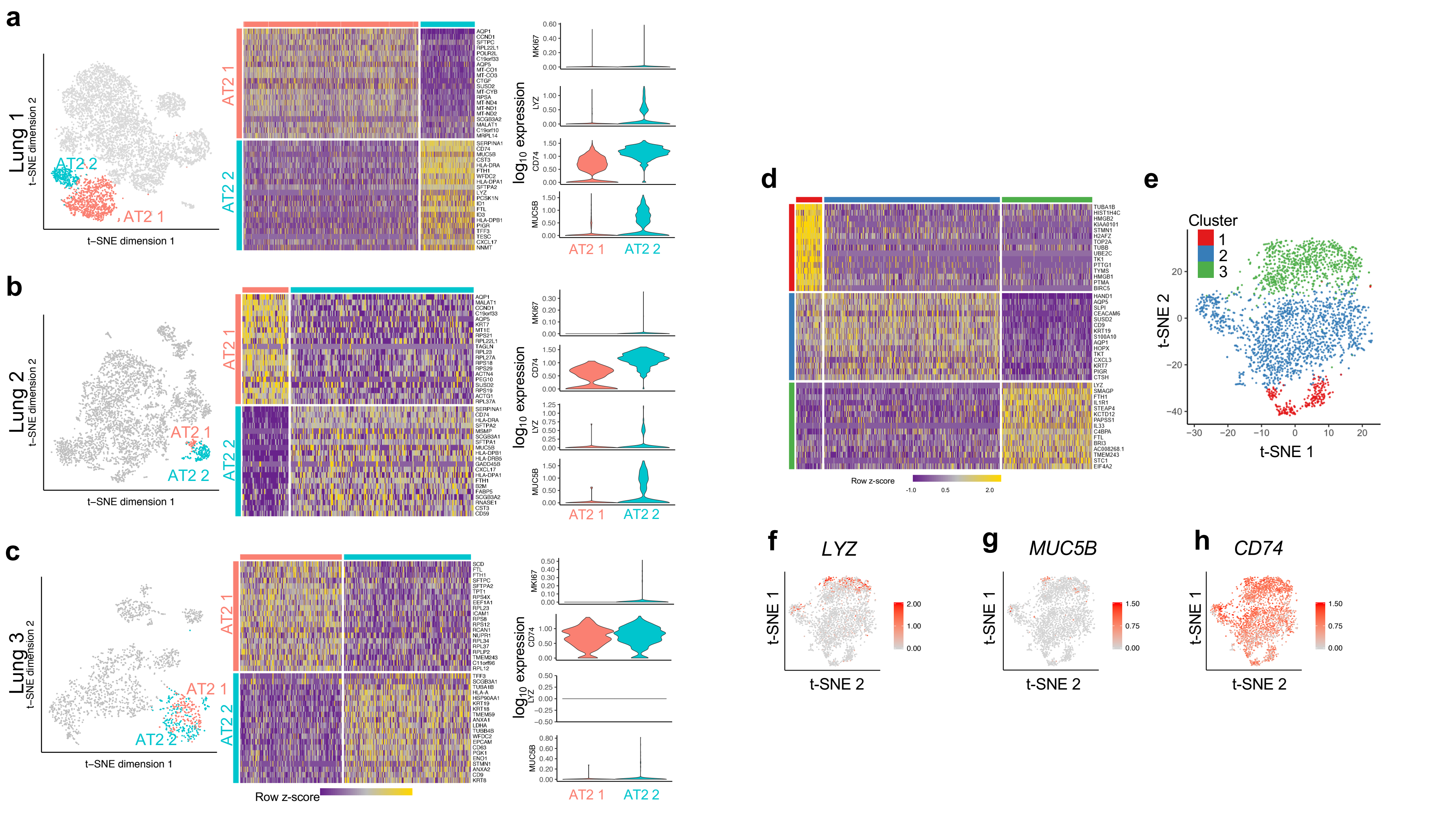


**Extended Data Figure 5.** Subclustering of AT2 cells. **a-c,** Upon configuring the parameter for the graph based cluster to create more clusters, AT2 populations were initially divided into two populations within the Lung 1 sample **(a)**. However, the same clustering procedure did not sub-divide the AT2 populations in the Lung 2 and Lung 3 samples **(b-c)**, likely due to fewer number of cells. Therefore, for these two samples in particular, we isolated the raw expression of the AT2 cells, and re-analyzed with Seurat and graph-based clustering was solely performed on these cells. The clustering parameter was set such that two clusters would be created and compared the population structure with that in Lung 1. **d-h,** scRNA-seq analysis of 2,780 cells in organoid culture derived from FACS purified AT2 cells after 89 days of organoid culture from Figure 2.. **d-e,** Clustering and t-SNE projection of AT2 cells demonstrates a proliferative subcluster (Cluster 1) that is defined by cell cycle genes per GSEA (**Supplementary Data 4**) but does not correspond to a specific AT2 subpopulation. **f-h,** Feature plots of *LYZ*, *MUC5B*, and *CD74* do not highlight discrete AT2 subpopulations that correspond to those seen in mixed organoid cultures (**Extended Data Fig. 4**), All plots are $\log_{10}$ expression.

### Extended Data Figure 6

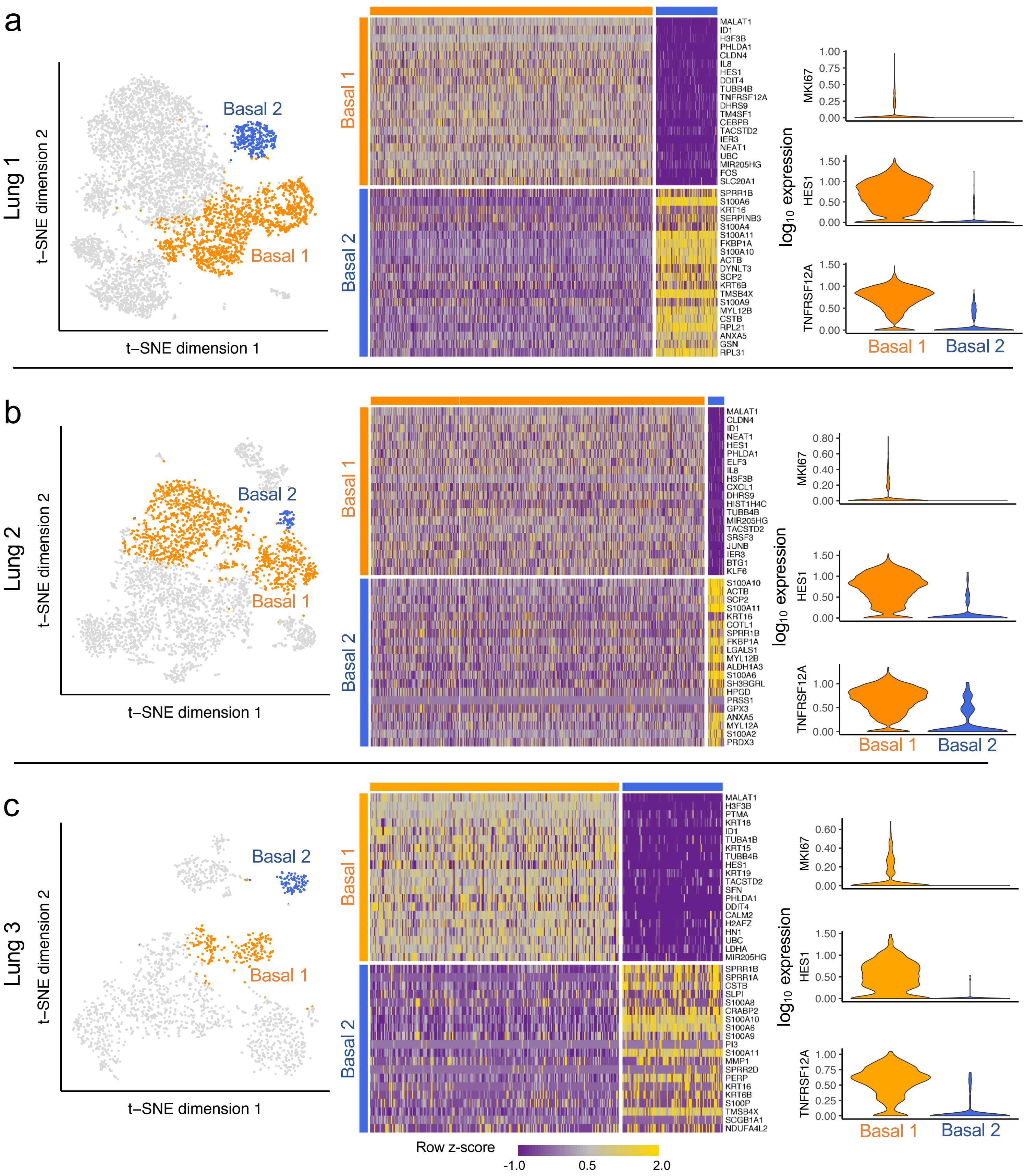


**Extended Data Figure 6.** scRNA-seq identifies an active basal cell subpopulation across three individual patient organoid cultures. **a-c,** High resolution clustering analysis identifies a reproducible active basal cell subpopulation with significantly higher expression of the surface marker *TNFRSF12A*, the NOTCH pathway marker *HES1*, and the proliferation marker *MKI67*. Modified Kruskal-Wallis Rank Sum Test p-values: *TNFRSF12A* 4.15 x $10^{-8}$; *HES1* 2.4 x $10^{-10}$; *MKI67* 3.4 x $10^{-3}$.

### Extended Data Figure 7

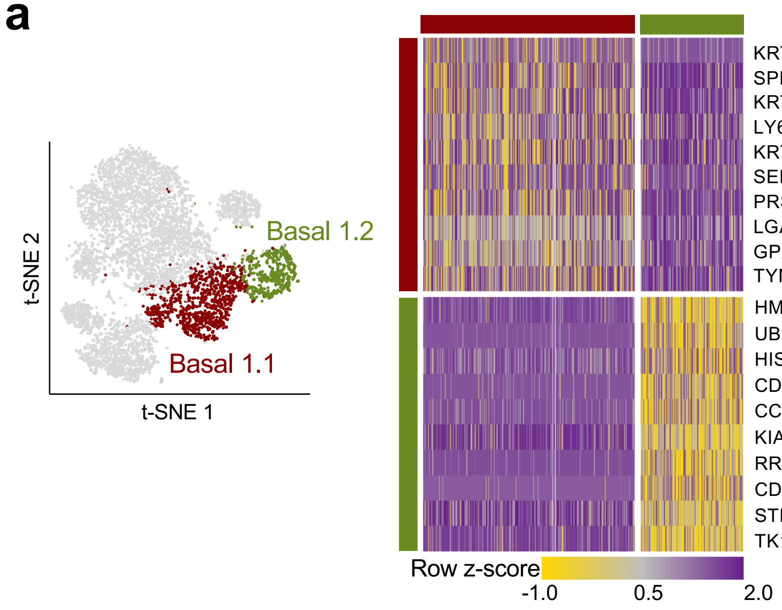


**b**

| GO Biological Process | p-value |
|---|---|
| cell cycle (GO:0007049)* | 3.90E-56 |
| cell cycle process (GO:0022402)* | 9.39E-52 |
| mitotic cell cycle (GO:0000278)* | 4.21E-46 |
| mitotic cell cycle process (GO:1903047)* | 2.38E-41 |
| chromosome organization (GO:0051276)* | 2.38E-39 |
| cell division (GO:0051301)* | 3.21E-35 |
| organelle organization (GO:0006996) | 5.25E-32 |
| DNA metabolic process (GO:0006259) | 3.95E-26 |
| cell cycle phase transition (GO:0044770)* | 1.69E-25 |
| chromosome segregation (GO:0007059)* | 4.35E-25 |
| nuclear chromosome segregation (GO:0098813)* | 6.20E-25 |
| mitotic cell cycle phase transition (GO:0044772)* | 1.03E-24 |
| sister chromatid segregation (GO:0000819)* | 3.15E-24 |
| nuclear division (GO:0000280)* | 1.03E-21 |
| DNA replication (GO:0006260)* | 2.44E-21 |
| DNA conformation change (GO:0071103) | 2.78E-21 |
| organelle fission (GO:0048285)* | 2.46E-20 |
| mitotic nuclear division (GO:0140014)* | 4.69E-20 |
| cellular component organization (GO:0016043) | 5.50E-20 |
| regulation of cell cycle (GO:0051726)* | 1.62E-19 |

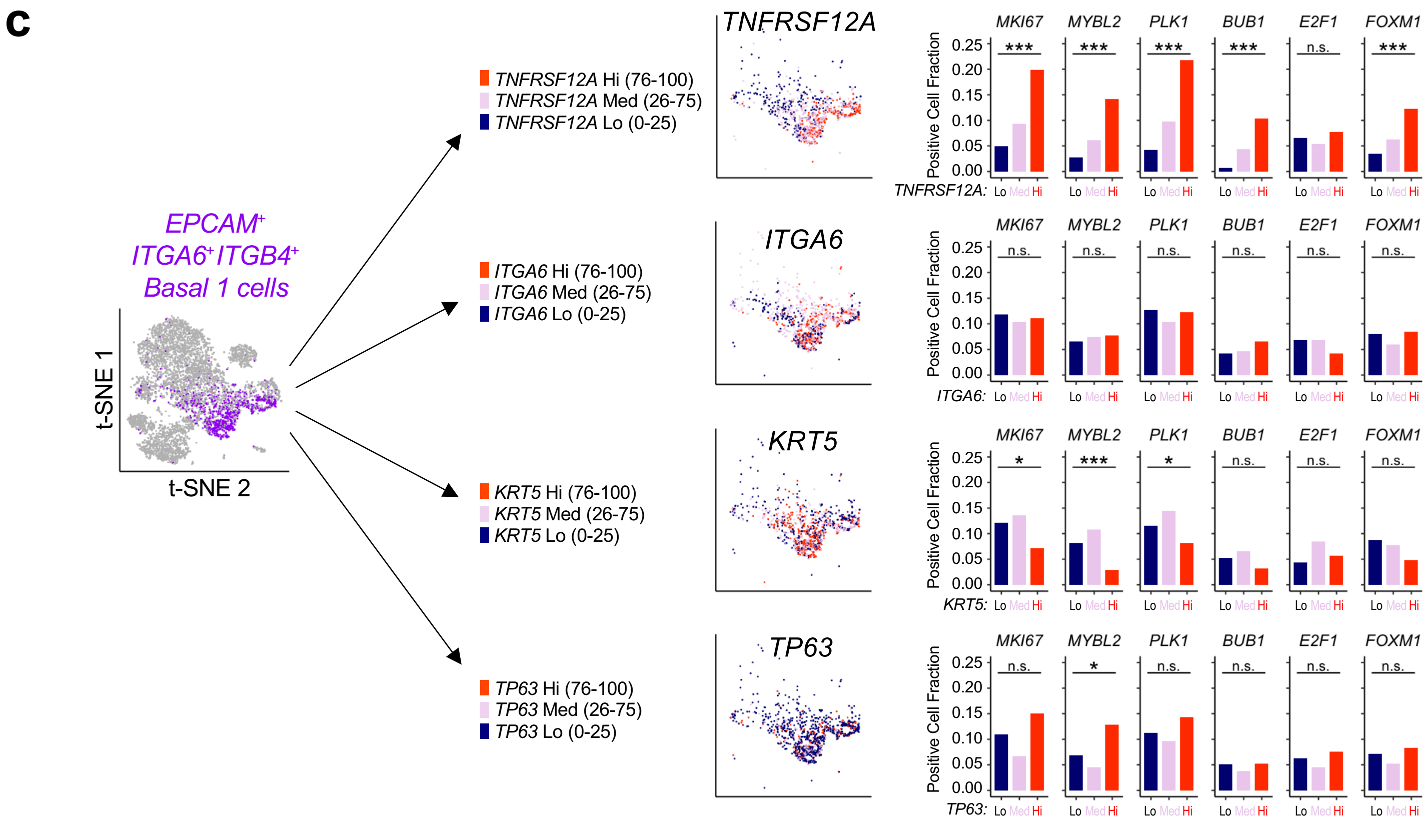


**Extended Data Figure 7.** scRNA-seq analysis of proliferation within the Basal 1 population **a,** Fine resolution clustering of *KRT5*+ populations identifies two Basal 1 sub-clusters, Basal 1.1 and 1.2. **b,** Gene Ontology PANTHER overrepresentation of differentially expressed genes enriched in Basal 1.2 versus 1.1 show the majority of Basal 1.2 processes involve cell cycle (asterisks). Complete analysis is provided in **Supplementary Data 4. c,** scRNA-seq subfractionation of cells simultaneously expressing EPCAM, ITGA6 and ITGB4 mRNA from Fig. 3d, according to highest quartile (Hi, 76-100%), medium (Med, 26-75%) or lowest quartile (Lo, 0-25%) mRNA expression of the indicated transcripts *(TNFRSF12A, ITGA6, KRT5* and *TP63*). Amongst the Lo, Med and Hi fractions of each Basal 1 marker, the cell fraction positive for the indicated cell cycle mRNAs *(MKI67, MYBL2, PLK1, BUB1, E2F1, FOXM1)* are plotted in the bar graphs. TNFRSF12A mRNA expression strongly enriches for cell cycle mRNAs over the progression from Lo→Med→Hi, which is not observed for ITGA6, KRT5 and TP63 mRNA expression does not. For *TNFRSF12A* and *ITGA6,* Hi, Med, and Lo correspond to the top 25%, mid 50%, and bottom 25% quartiles respectively. However, for *KRT5* and *TP63*, a significant number of cells expressed 0 UMI, so we defined Lo as cells with zero expression, and Hi and Med as the top 50% and bottom 50% of cells with non-zero expression. Thus, *KRT5* has Hi (33.7%), Med (33.7%), and Lo (32.6%), and *TP63* has Hi (12.8%), Med (12.8%), and Lo (74.4%).

### Extended Data Figure 8

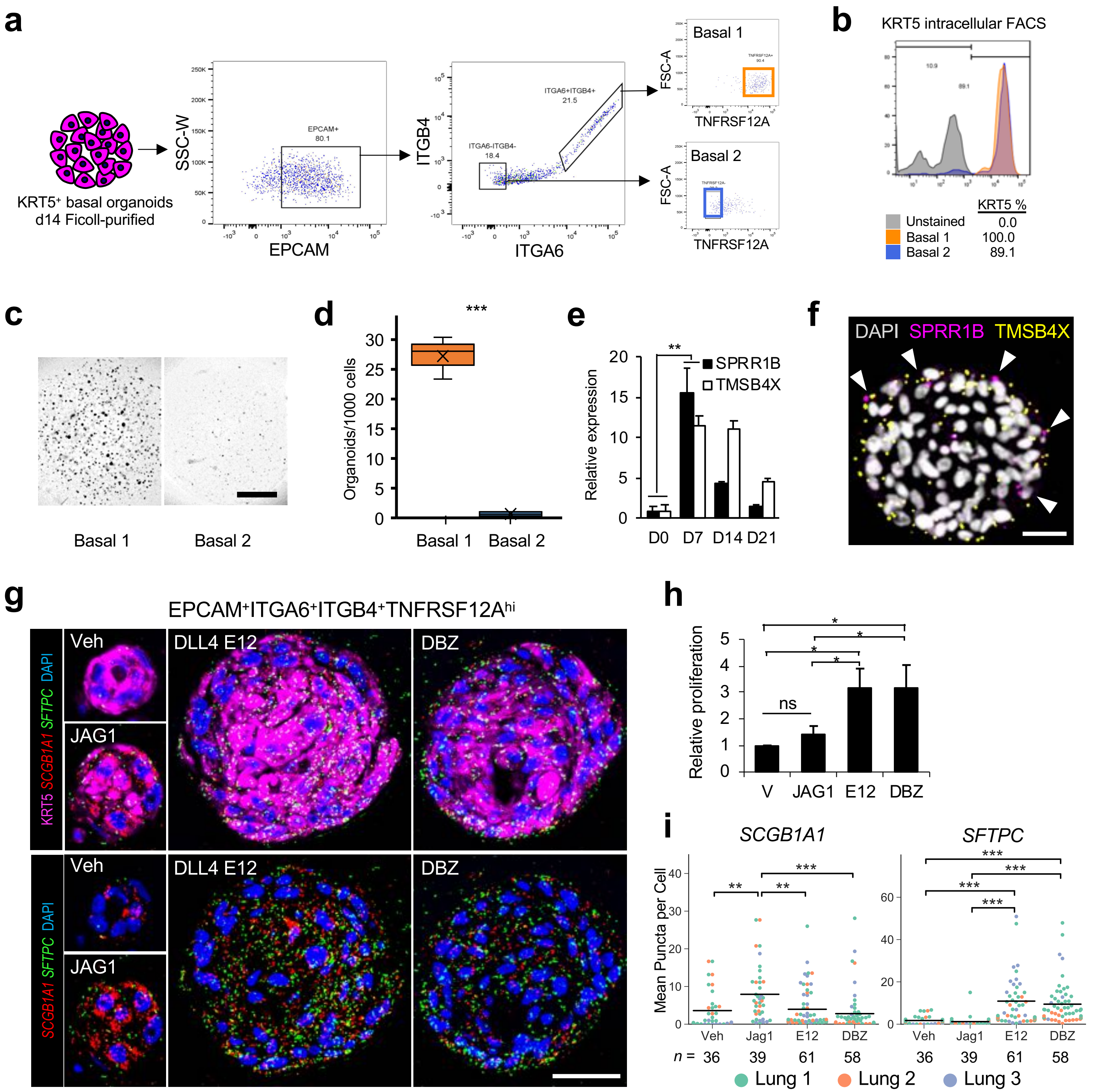


**Extended Data Figure 8.** Evaluation of Basal 1 lineage relationship to Basal 2 and the influence of NOTCH signaling on Basal 1 renewal and differentiation. **a,** Isolation of Basal 1 and Basal 2 via differential sedimentation of KRT5⁺ cells followed by FACS sorting of EPCAM⁺ITGA6⁺ITGB4⁺TNFRSF12A⁺ (Basal 1) versus EPCAM⁻ITGA6⁻ITGB4⁻TNFRSF12A⁻ (Basal 2). **b,** Intracellular FACS measurement of KRT5⁺ protein expression in Basal 1 and 2 fractions from (a). **c**, representative brightfield of day 14 cultures from (a-b). **d,** quantitation of 3 biologic replicates from (a-c) (*** $p < 0.001$ two tailed t-test). **e,** qPCR measurement of two differentially upregulated Basal 2 genes from the three scRNA-seq biological replicates (**Extended Data Fig. 6**, *SPRR1B*, *TMSB4X*) after prolonged culture of FACS isolated Basal 1 cells. Data are relative mean +/- SEM of cultures from three individuals, ** = $p < 0.01$. **f,** RNA FISH demonstrating *TMSB4X* and *SPRR1B* cellular transcripts within organoids originating from Basal 1 cells (arrows), scale bar = 25 µm. **g,** KRT5 immunostaining and *SFTPC* and *SCGB1A1* RNA FISH of FACS isolated TNFRSF12A$^{hi}$ Basal 1 cells under vehicle, NOTCH agonism (JAG1 peptide), or NOTCH antagonism with the Delta-like ligand mutant 4 (DLL4$^{E12}$) or the gamma secretase inhibitor DBZ; scale bar = 50 µm. **h,** Fluorescent quantitation of resazurin dye reduction to estimate relative cellular proliferation in (a), data represent mean and SEM of cultures from five individuals, * $p < 0.05$ two-tailed Student's t-test. **i,** Quantitation of *SCGB1A1* and *SFTPC* gene expression by RNA FISH in the context of NOTCH agonism or antagonism, ** $p < 0.01$, *** $p < 0.001$ Student's t-test.

### Extended Data Figure 9

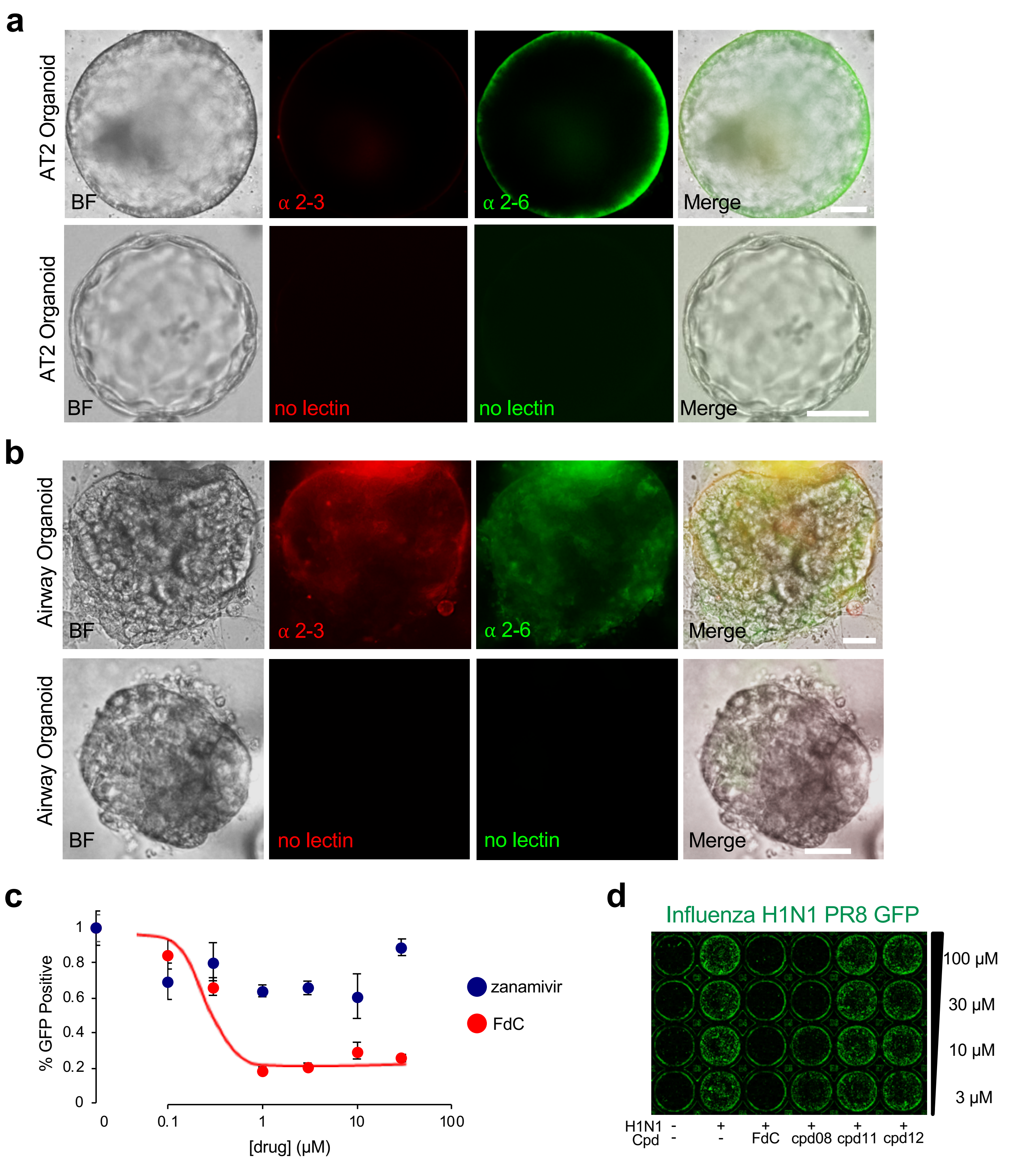


**Extended Data Figure 9.** Characterization of mixed distal lung organoid culture for influenza infection modeling. **a-b,** Lectin staining with *M. amurensis* (α 2-3) and *S. nigra* (α 2-6) lectins or no lectin negative controls to characterize sialic acid residues which serve as surface molecules for influenza virus host cell entry. Scale bar = 25 μm. **c,** Dose response curves for two different classes of antiviral drugs on influenza infectivity and replication. As expected, the nucleoside analog FdC demonstrated an $IC_{50}$ of 340 nM as compared to zanamivir, which did not exhibit a dose response curve and an $IC_{50}$, a neuraminidase antagonist which only impairs viral shedding, but not infectivity and replication. N = 3 technical replicates. **d,** Fluorescence micrograph of multiwell screening of selected various antiviral agents after H1N1 PR8-GFP organoid infection in 48 well format. FdC= nucleoside analog 2'-deoxy-2'-fluorocytidine. Cpd= compound #.

### Extended Data Figure 10

#### Eversion and accelerated ciliary differentiation of apical-out basal organoids

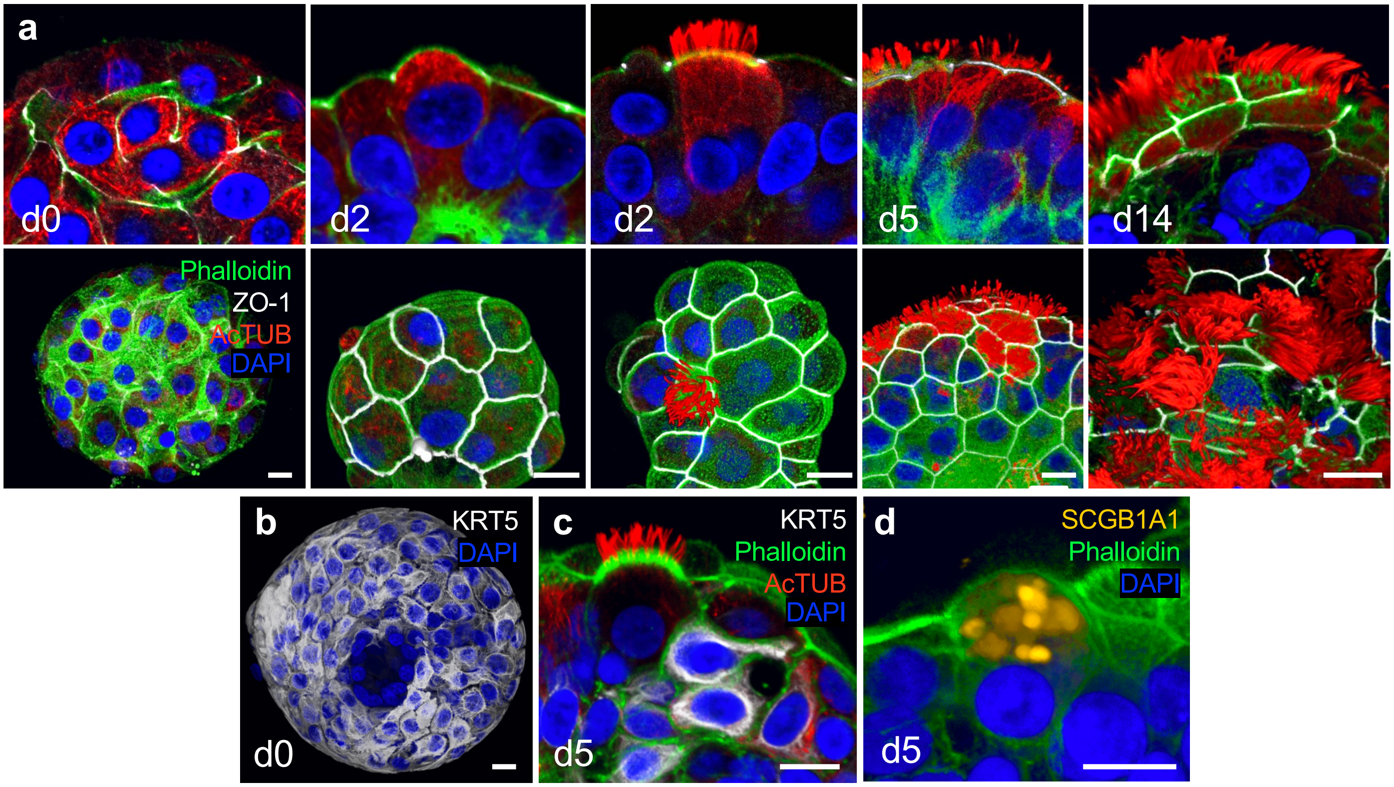


#### Prolonged suspension culture of AT2 organoids induces apical-out polarization and AT1 differentiation

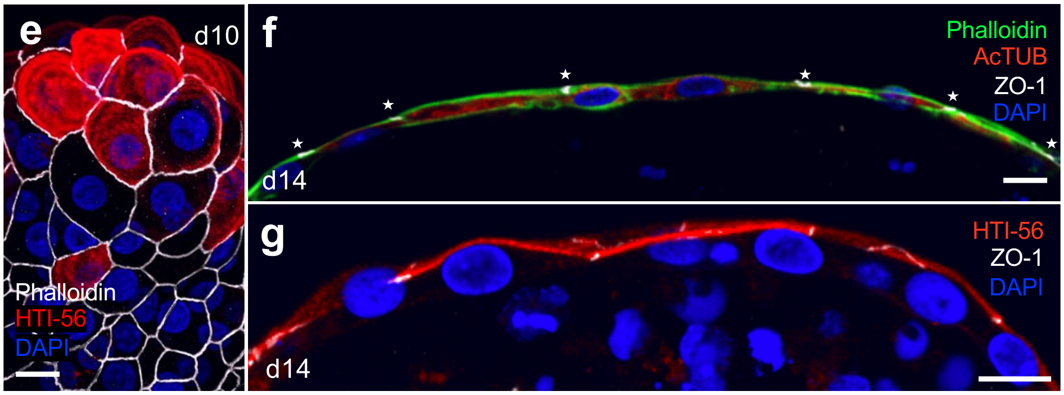


**Extended Data Figure 10. Apical-out polarization and multi-lineage differentiation of distal lung organoids upon suspension culture.** **a-d,** Eversion and accelerated ciliary differentiation of apical-out basal organoids. **a,** Confocal 3D sections (top panels) and surface reconstructions (bottom panels) of apical-out lung organoids at different days after ECM removal. At day 0 (d0) microfilament (green, phalloidin) and microtubule (red, acetylated tubulin) organization is not polarized. Junctional strands (ZO-1, white) are discontinuous and found between cells inside of organoids. By day 2 in suspension (d2) ZO-1 (white) forms junctional rings in the apical periphery of each cell facing the external side of the organoids and the actin cytoskeleton forms microvilli (green) facing outward (apical-out polarity). Also at d2 some cells initiate microtubule polarization and formation of tufts of cilia. By live-cell imaging (see Supplemental Video 3) cilia become motile by day 5. By day 5 (d5) many more cells have differentiated into multiciliated respiratory epithelium with motile cilia facing outwards. Mature motile cilia can be observed for several weeks, example at day 14 (d14), (Supplemental Video 3). **b,** 3D confocal reconstruction of an organoid embedded in ECM consisting mostly of basal stem cells (KRT5$^+$, white). **c,** As apical-out polarity is established and ciliagenesis begins, KRT5$^+$ cells decrease in abundance. Basal cells are found underneath the polarized epithelium. **d,** SCGB1A1$^+$ Club cells with apical-out polarity can are present on the exterior everted surface. In all panels nuclei are stained blue with DAPI, and actin microfilament organization visualized with phalloidin (green). Scale bars = 10 µm. **e-g,** Prolonged suspension culture of AT2 organoids (day 10 post-eversion) induces apical-out polarization and AT1 differentiation. **e,** Optical sections through alveolar-derived organoids after 10 days in suspension culture show decreased abundance of AT2 cells while individual cuboidal cells begin to express the AT1 marker HT1-56 (red), a transmembrane protein specific to the apical membrane of alveolar type 1 pneumocytes (AT1). **f-g,** Side views of alveolar organoids after 10 days of suspension culture reveal thin AT1 cells with apical junctional complexes facing outwards (apical-out) and expression of HT1-56 on the apical membrane.
