## Supplemental_Files for "Progenitor identification and SARS-CoV-2 infection in long-term human distal lung organoid cultures": 315402_5_supp_3462051_qcf2k4.docx

**Supplementary Information Guide**

1. **Supplementary Methods**
2. **Supplementary Tables**

Supplementary Table 1 – Clinical demographics of 103 individuals from whom normal lung was obtained.

Supplementary Table 2 – Basal 1 membrane proteins (Identified by ontology analysis of top differentially expressed genes with GO term 0031224, intrinsic component of membrane).

Supplementary Table 3 – Next Generation Sequencing of five distal lung organoid cultures. Summary and annotations of nonsynonymous variants with allele frequency > 0.10.

3. **Supplementary Data**

Supplementary Data 1 – genes used for SPADE analysis

Supplementary Data 2 – FACS gating strategies

Supplementary Data 3 – scRNA-seq analysis in R markdown html format

Supplementary Data 4 – Gene Set Enrichment Analysis (GSEA) of top differentially expressed genes from populations corresponding to Basal 1, Basal 2, Basal 1.1, Basal 1.2, and Cluster 1 of purified AT2 as well as corresponding top differentially expressed genes for each population. Bold = genes used for GSEA analysis.

Supplementary Data 5 – High resolution confocal micrographs corresponding to Z projections in Fig. 5a.

Supplementary Data 6 – Variants identified by Next Generation Sequencing within five distal lung organoid cultures (TOMA COMPASS Tumor Mutational Profiling System) in Variant Call Format.

Supplementary Data 7 – Quantitation and statistical analysis of SARS-CoV-2-infection among KRT5, SCGB1A1, and AcTUB cells.

4. **Supplementary Video**

Supplementary Video 1 – Transmission confocal microscopy video of beating cilia (25X magnification).

Supplementary Video 2 – Bright-field microscopy video of beating cilia (10x magnification).

Supplementary Video 3 – DIC and 3D confocal microscopy video of motile cilia in apical-out basal organoids at suspension culture day 5 and 14.
