## Supplemental_Files for "Progenitor identification and SARS-CoV-2 infection in long-term human distal lung organoid cultures": 315402_5_source_data_3462038_qcfwff.pdf

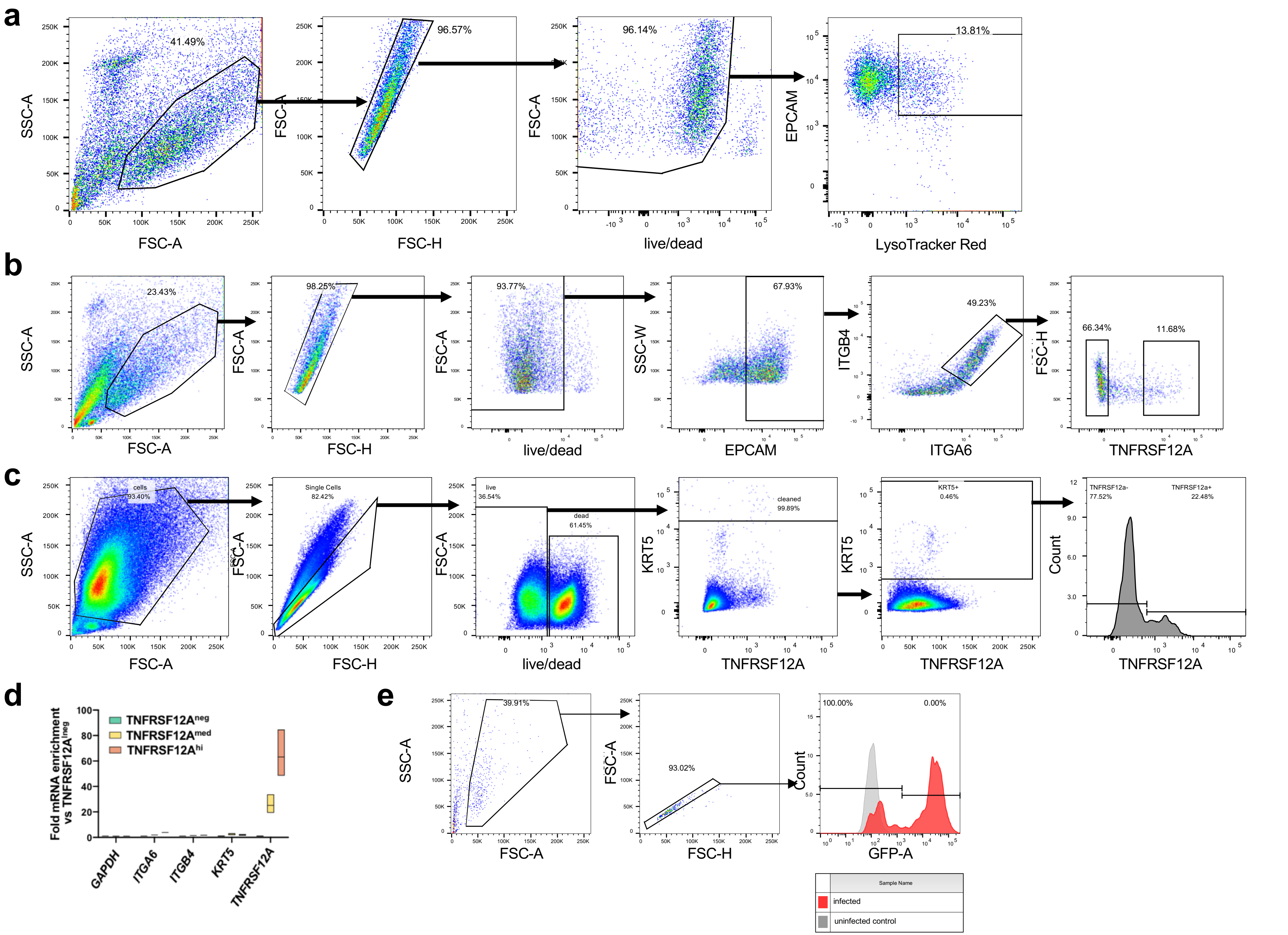


Supplementary Data 2. Gating strategies. **a**, EPCAM$^{+}$Lysotracker$^{+}$ gating strategy to purify AT2 cells. Purity was confirmed with 100/100 sorted cells staining positive for SFTPC protein. **b**, EPCAM$^{+}$ITGA6$^{+}$ITGB4$^{+}$TNFRSF12A$^{hi/lo}$ gating strategy. Purity was confirmed with 100/100 pooled sorted cells staining positive for KRT5 protein. **c**, Gating strategy to analyze KRT5+ cells and corresponding TNFRSF12A$^{+}$ populations within cells freshly dissociated from distal human lungs. **d,** RT-PCR correlation of TNFRSF12A and other mRNAs from cells sorted by FACS into TNFRSF12A neg, med, and hi fractions **e**, Gating strategy to estimate H1N1 PR8 GFP infectivity in human lung organoids.
