## Supplemental_Files for "Progenitor identification and SARS-CoV-2 infection in long-term human distal lung organoid cultures": 315402_5_supp_3462050_qcfznc.pdf

### Supplementary Methods

### Lung Organoid Media Components

| Component | Vendor | Cat # | [Final] |
|---|---|---|---|
| HEPES | Gibco | 15630-080 | 1 mM |
| GlutaMAX-1 (100x) | Gibco | 35050-061 | |
| NIC (Nicotinamide) | Sigma | N0636 | 10 mM |
| NAC (N-acetylcysteine) | Sigma | A9165-5G | 1 mM |
| B-27 supplement (50x) | Gibco | 125870-01 | 1X |
| A83-01 | Tocris | 2939 | 500nM |
| Pen/Strep Glutamine | Gibco | 10378016 | |
| Human EGF | R&D | 236-EG-01M | 50 ng/mL |
| Human Noggin | R&D | 6057-NG/CF | 100 ng/mL |
| Advanced DMEM/F-12 | Thermo Fisher | 12634-028 | |

### Additional Medium Additives

| Component | Vendor | Cat # | [Final] |
|---|---|---|---|
| Wnt-3a | R&D | 5036-WN | 100ng/mL |
| R-spondin 1 | Peprotech | 120-38 | 500ng/mL |
| WNT C-59 | Biogems | 1248913 | 1μM |

### Immuno- and Lectin- fluorescence Reagents

| Anti-body | Vendor | Catalog # | Dilution |
|---|---|---|---|
| Mouse anti-HTII-280 | Terrace Biotech | TB-27AHT2-280 | 1:100 |
| Mouse anti-HTI-56 | Terrace Biotech | TB-29AHT1-56 | 1:100 |
| Rabbit anti-SFTPC | Santa Cruz Biotechnology | sc-13979 | 1:100 |
| Mouse anti-Acet. Tubulin | Sigma Aldrich | T7451 | 1:100 |
| Rabbit anti-Cytokeratin 5 | AbCam | Ab193895 | 1:400 |
| Mouse anti-SCGB1A1 | Santa Cruz | Sc-365992 | 1:100 |
| Goat anti-ACE2 | AbCam | AF933 | 1:100 |

| | | | |
|---|---|---|---|
| Sheep anti-SCGB1A1 | Novus | AF4218 | 1:100 |
| Rabbit anti-ZO-1 | Invitrogen | 617300 | 1:100 |
| Mouse anti-SARS-CoV2 NP | Sinobiological | 40143-MM05 | 1:100 |
| Mouse anti-dsRNA | Scicons | 10010200 | 1:100 |
| Rabbit anti-acetyl-α-tubulin | Cell Signaling | 5335 | 1:100 |
| Blocking Serum | Jackson Immunoresearch | 017-000-121<br>005-000-121 | 1:10 |
| Secondary Antibodies | Jackson Immunoresearch | 711-165-152<br>711-095-152<br>715-165-151<br>715-095-151 | 1:400 |
| Anti-mouse IgM | Invitrogen | A-21042 | 1:400 |
| Anti-mouse IgG | Invitrogen | A-21422 | 1:400 |
| Alexa Fluor 660 Phalloidin | Thermo Fisher | A22285 | 1:100 |
| Mouse anti-TNFRSF12A | Biolegend | Clone ITEM-4 314102 | 1:100 |
| Rabbit anti-TNFRSF12A | Thermo Fisher | PA5-20275, lot TD2561104 | 1:100 |

**qRT-PCR Reagents**

| | Reagent | Vendor | Catalog # |
|---|---|---|---|
| RNA Isolation | PicoPure™ RNA Isolation Kit | Applied Biosystems | KIT0214 |
| cDNA Synthesis | iScript™ cDNA Synthesis Kit | Bio-Rad | 1708841 |
| Pre-amplification | SsoAdvanced™ PreAmp Supermix | Bio-Rad | 1725160 |
| Taqman Gene Expression Assay | *GAPDH* | Thermo Fisher | Hs02786624_g1 |
| | *ITGA6* | Thermo Fisher | Hs01041011_m1 |
| | *ITGB4* | Thermo Fisher | Hs00236216_m1 |
| | *KRT5* | Thermo Fisher | Hs00361185_m1 |
| | *TNFRSF12A* | Thermo Fisher | Hs00171993_m1 |

| | | | |
|---|---|---|---|
| COV2 RT-PCR Assay | nCOV_N1 FWD | IDT | 10006821 |
| | nCOV_N1 REV | IDT | 10006822 |
| | nCOV_N1 Probe | IDT | 10006823 |
| | TaqMan™ Universal Master Mix II, no UNG | Applied Biosystems | 4440040 |

qRT-PCR Primers

| | Sequence 5' → 3' |
|---|---|
| SARS unspliced FWD | TGACTTCACGGAAGAGAGGTT |
| SARS unspliced REV | AACAGTTAACACAATTTGGGTGG |
| Human U3 FWD | CGTGTAGAGCACCGAAAACC |
| Human U3 REV | AACAGTTAACACAATTTGGGTGG |

**Computational and Statistical Analysis of the scRNA-seq Data**

scRNA-seq analysis was initially carried out on the data set with the greatest number of cells (Lung 1, 7,285 cells), and the same analysis were repeated on Lung 2 (4,512 cells) and Lung 3 (3,364 cells).

**Data Preprocessing**

Sample multiplexing, barcode processing, RNA-seq alignment, and 3' gene counting were performed with the Cell Ranger Software Suite[1] (version 1.1). We used Seurat[2], an R toolkit developed for single cell analysis, to perform basic quality control of the data. Cells with less than 200 UMIs were filtered out, and genes expressed in less than 3 cells were filtered out. To further improve the quality of the data, we used the workflow recommended by Seurat on droplet-based scRNA-seq data (http://satijalab.org/seurat/pbmc-tutorial.html) to regress out (i) the effects of the total number of UMIs and (ii) the percentage mitochondrial gene counts. In addition, as suggested by the workflow, we used the gene dispersion analysis[3] implemented in Seurat to select highly variable genes, retaining genes with logarithmic (base 10) mean expression between 0.05 and 5 and with logarithmic dispersion greater than 0.2. The quality control resulted in 7285 single cells and 1938 highly variable genes for unbiased analysis.

**Unbiased Clustering and Visualization**

First, we applied principle component analysis (PCA) to reduce the dimension of the data. The implementation of PCA in Seurat[2] scaled the data to zero mean and unit variance on the log-transformed data. The principle components (PCs) were sorted by their ability to explain the variance in the gene expression matrix. In order to select the number of PCs, we used the jackstraw procedure[4], which is a built-in boot-strap procedure implemented in Seurat that resamples 1% of the data, re-runs PCA, and scores each PC accordingly. As part of the jackstraw procedure, the p-value for each PC is computed by a proportion test comparing the number of genes with a p-value (testing the significance of association between a gene with the PC) below a particular threshold (default given by $10^{-5}$), compared with the number of genes expected under a uniform distribution of the gene p-values. We selected the top 23 PCs based on the increase from $10^{-23}$ to $10^{-19}$. We did not notice significant difference downstream with more PCs as they did not seem to capture additional signals we could interpret.

We applied unsupervised clustering to partition cell populations into groups to identify meaningful structures in the data. In particular, we applied a graph based clustering technique on the top 23 PCs (also implemented in Seurat), tailored for high-dimensional single-cell data. The method embeds cells in a K-nearest neighbor (KNN) graph and partitions the graph into highly interconnected cell communities[5]. The clustering algorithm requires a resolution parameter that determines the number of distinct cell populations. Varying the parameter from 0.1 to 0.5 yields 5 to 11 clusters representing cell identities defined at different resolutions.

To visualize the data, we used the t-Stochastic Neighbor Embedding (t-SNE) algorithm[6], as often done with scRNA-seq data, given its ability to capture non-linear relationships. We applied t-SNE on the significant PCs to visualize the cells in 2-D. In the t-SNE 2-D mapping (also referred to as the t-SNE space), each point corresponds to a cell, and cells that share similar global gene expression appear closer together in the 2-D map and cells that are different appear further apart. Furthermore, t-SNE enables us to visualize the expression levels of each cell for given genes, as well as groupings by technical batches or by clustering results.

### Major Cell-type Annotation

The clustering algorithm is tunable and can result in grouping at different resolutions, or simply different number of groups. Therefore, we considered the clusters provided at a low resolution and a high resolution to determine the appropriate number of major cell types. In order to identify the major cell types that are present among the clusters, we performed differential gene analysis to identify cluster specific gene markers. We enumerated genes that are most up regulated in a given population relative to all the other populations. In particular, we computed the average of the log10 expression of each gene among the cells assigned to a given cluster, and compared this average value to the average log10 expression of the remaining cells in the data after normalization and size correction implemented in sSeq[7]. After analyzing the top genes for each high-resolution cluster, we merged clusters that shared highly expressed genes and annotated 5 major populations, including basal cells, ATII cells and club cells.

### Testing Canonical Marker Genes

Based on our experimental setup and staining experiments, we expected at least three populations (ATII, club and basal) to be present in the data. We wanted to see if the canonical markers for these cell types are significantly differentially expressed across the populations. However, because the expression levels of these same genes were part of the data used to identify cell clusters, naive tests comparing the expression levels of these markers across clusters are not valid.

To work around this issue, we studied how the clusters we identify are dependent on the canonical cell type markers. First, we repeated the unsupervised clustering analyses using the exact same procedure described above on a scRNA-seq data excluding measurements for each of the marker genes. This yields clustering assignments that are "independent" of the marker gene (given the correlation that exists among the expression of many genes, we are not referring here to stochastic independence). We compared the low-resolution clusters (parameter 0.1 for the graph based clustering) obtained with and without each of the marker genes. After verifying that the clustering results were highly consistent according to multiple clustering concordant metrics, we used Kruskal-Wallis Rank Sum Test to test if a known marker gene is differentially expressed across all populations based on the clusters that did not utilize information of the marker gene (with the null hypothesis that the mean expression of a given gene is the same across all groups.)

### Exploratory Analysis of Basal Cells

We found the basal cell population to be highly heterogeneous, so we analyzed the genes that appeared to be differentially expressed among the subpopulations. First, we identified from both t-SNE and clustering two distinct clusters that corresponded to a subpopulation of quiescent basal cells and a subpopulation of non-quiescent basal cells. To identify signatures of novel cell populations, we also searched for enriched markers that are surface markers. We intersected the most up-regulated or down-regulated genes with genes annotated with surface marker proteins (GO: 0031224, intrinsic component of membrane, **Supplementary Table 2**). Among the identified genes, we found the gene *TNFRSF12A* to be highly differentially expressed and one that would be amenable to FACS purification. In addition, we noticed *HES1* was also ranked as one of the top genes differentiating the non-quiescent basal population from the quiescent basal population.

Because *TNFRSF12A* and *HES1* were identified by ranking differentially expressed genes in only one sample, we conducted the exact same pipeline on two independent biological samples, where we also easily identified the non-quiescent and quiescent basal cells. We took the gene rankings for the three samples and applied robust rank aggregation[8] for the rank of each gene among the list. For each gene, the algorithm looks at how the gene is ranked in each list and compares this with the rankings to the baseline case where all the gene lists are randomly shuffled. As a result, each gene can be assigned a p-value, which we report as the significance of its rank.

### Associating TNFRSF12A with Proliferative Markers among Basal Cells

To understand the role of *TNFRSF12A*, we investigated associations between expression levels of *TNFRSF12A* and known proliferative markers both experimentally and computationally. Experimentally, the canonical basal cells were FACS sorted by the simultaneous presence of ITGA6, ITGB4 and EPCAM, followed by separating this triple-positive population into TNFRSF12A$^{neg}$ and TNFRSF12A$^{hi}$ fractions which were independently cultured to measure cell growth. To computationally mimic this approach and explore its outcome we selected all cells in scRNA-seq that expressed basal markers, and divided the cells into three groups defined by the top (*TNFRSF12A*$^{hi}$), bottom (*TNFRSF12A*$^{lo}$) and middle two (*TNFRSF12A*$^{med}$) quartiles of *TNFRSF12A* mRNA expression. Finally, we tested if the presence of a proliferative gene signature[9] (*MKI67, MYBL2, PLK1, BUB1, E2F1, FOXM1*) was associated with the these three groups using a Chi-square test for independence.

**Computational Reproducibility**

We provide a step-by-step guide on the scRNA-seq analysis used for this manuscript in R markdown html format using the R knitr package[10]. The data objects used for the analysis are also included as **Supplementary Data 2**, with which one can reproduce the entire computational workflow of the scRNA-seq analysis.

**SPADE Analysis**

To explore and visualize the differentiation hierarchy between the cells in the data, we used a similar approach[11,12] based on SPADE (Spanning Tree Progression of Density-normalized events)[11] which uses a minimum spanning tree (MST) to represent the hierarchical relationship between cells. Cells within or in nearby branches on the tree are hierarchically more related compared to distant cells. The tree was optimized from a set of 42 genes comprised of a mixture of 15 highly expressed cluster-specific genes derived from the top 3 genes enriched in each of the 5 major t-SNE clusters described above in addition to 27 high variable genes derived using the robust statistics of variability known as median absolute deviation (MAD)[13] defined as median of the absolute deviations from the data's median (**Supplementary Data 1**). For each gene, the MAD score is estimated from the non-zero count data using R. The 27 high variable genes were derived from combining top genes with high MAD scores from cells within and between clusters using a supervised approach.

**Running SPADE**

SPADE analysis was performed using the R implementation[11], that runs on Mac OS X and requires an FCS data file input format created from asinh-transformed gene expression counts with cofactor of 5. The SPADE analysis involves running the following 4 major steps: (1) density-dependent down-sampling of single-cell data, (2) agglomerative clustering, (3) joining clusters with a minimum spanning tree and (4) up-sampling to map all cells onto the final output tree. Each node and node size in the MST output represents a median marker expression and the number of cells within that node respectively. SPADE was run using the default settings on an initial number of 200 clusters but with no density dependent down-sampling. To investigate the sensitivity of the tree layout, we performed 3 different random seeded SPADE analyses on the data set and visually compared the SPADE trees.

**SPADE Analysis of Club and Basal cells**

To investigate in more detail the hierarchical relationship between Basal and Club cells alone, a SPADE tree derived from using the same set of 42 genes described above was applied. The result is summarized in **Extended Data Fig. 3** which suggests potential intermediate states (outlined in purple) taking place in addition to the main Basal to Club transition (continuum from blue to red states) identified by visualizing the median expression profiles (**Extended Data Fig. 3**) for some major cluster specific and transition genes on the SPADE tree.

#### Next Generation Sequencing of Organoid Cultures

To verify the absence of cancerous cells in organoid culture, genomic DNA was extracted from five long-term organoid cultures (range 4-8 weeks) from different individuals and sequenced using a commercial targeted resequencing assay for 130 cancer genes and software (TOMA tumor profiling system, Foster City, CA). COSMIC annotated Variants are summarized in **Supplementary Table 3** and Variant Call Files are provided in **Supplementary Data 6**.

#### Organoid RNA Extraction, cDNA Synthesis, and qRT-PCR

Mixed human distal lung organoids (p0, day 28) were FACS sorted into TNFRSF12A-negative, -medium and -high fractions. RNA was extracted using the Arcturus PicoPure kit (Applied Biosystems, KIT0204). Extracted RNA was reverse-transcribed to cDNA using the iScript cDNA synthesis kit (Bio-Rad,1708841). SsoAdvanced PreAmp Supermix (Bio-Rad, 1725160) was used for pre-amplification. The following Taqman Gene Expression Assay were used (Thermo Fisher): GAPDH as endogenous control (Hs02786624_g1), Krt5 (Hs00361185_m1), ITGA6 (Hs01041011_m1), ITGB4 (Hs00236216_m1), TNFRSF12A (Hs00171993_m1) along with TaqMan™ Universal Master Mix II, no UNG (Applied Biosystems, 4440040). All samples were run in triplicate on Applied Biosystems 7900HT Fast Real-Time PCR System with cycle conditions: 50C for 2 min, 95C for 10 min, followed by 40 cycles of 95C for 15 sec, and 60C for 1 min. Fold change of gene expression of TNFRSF12A-negative, -medium and -high fractions compared to the TNFRSF12A-negative sample was calculated using the ΔΔCt method and presented in **Supplementary Data 2**.

#### RNA FISH using Proximity Ligation- in situ Hybridization (PLISH)[14]

RNA FISH on paraffin embedded sections was performed according to the protocol by Nagendran et al. for *SFTPC* and *SCGB1A1* with the corresponding probes below:

| | Probe Sequence 5’ → 3’ |
|---|---|
| ***SFTPC*** | |
| SFTPC-HL2X | TCGTACGTCTAACTTACGTCGTTATGTTGTAGGCGATCAGCAGCTG |
| SFTPC-HR2X | AGCAGGTGCCAGGGGCTGGCTTATACGTCGAGTTGAAGAACAACCTG |
| ***SCGB1A1*** | |
| SCGB1A1a-TxR 91 | TTAGTAGGCGAACTTACGTCGTTATGTGCAGCAGAGAGCCAGTGTG |
| SCGB1A1a-TxR 112 | GATCTCTGCAGAAGCGGAGCTTATACGTCGAGTTGAACATAAGTGCG |
| SCGB1A1b-TxR 166 | TTAGTAGGCGAACTTACGTCGTTATGAACTGGAGGGTGTGTCCATG |
| SCGB1A1b-TxR 187 | AAGTTCCATGGCAGCCTCATTTATACGTCGAGTTGAACATAAGTGCG |
| SCGB1A1c-TxR 278 | TTAGTAGGCGAACTTACGTCGTTATGTTAATGATGCTTTCTCTGGG |
| SCGB1A1c-TxR 299 | GGGCTATTTTTTCCATGAGCTTATACGTCGAGTTGAACATAAGTGCG |
| SCGB1A1d-TxR 374 | TTAGTAGGCGAACTTACGTCGTTATGTCAAAGCATGGCAGCGGCAG |
| SCGB1A1d-TxR 395 | CAAGGCTGGTGGGCGTGGACTTATACGTCGAGTTGAACATAAGTGCG |

| | |
|---|---|
| ***SPRR1B*** | |
| SPRR1B-HL3X | TATTCGTTCGAACTTACGTCGTTATGTATAAGGGAGCTGAATCATT |
| SPRR1B-HR3X | GAAAGTGAATTTAATGGGGGTTATACGTCGAGTTGACCGACGTATTG |
| ***TMSB4X*** | |
| TMSB4X-HL5X-57 | TAGCGCTAACAACTTACGTCGTTATGTTGCGGAGGAAAAGCGAAGC |
| TMSB4X-HR5X-57 | ATCGGGTTTGTCAGACATGGTTATACGTCGAGTTGAACGTCGTAACA |
| TMSB4X-HL5X-71 | TAGCGCTAACAACTTACGTCGTTATGTTTGTCAGACATGGTTGCGG |
| TMSB4X-HR5X-71 | TCGATCTCAGCCATATCGGGTTATACGTCGAGTTGAACGTCGTAACA |
| TMSB4X-HL5X-477 | TAGCGCTAACAACTTACGTCGTTATGTTTTACTCTAGATTTCACTG |
| TMSB4X-HR5X-477 | GACACCTTGGGCCAGCTTGGTTATACGTCGAGTTGAACGTCGTAACA |
| TMSB4X-HL5X-9 | TAGCGCTAACAACTTACGTCGTTATGTCTGCGCAGTGGCCACCACC |
| TMSB4X-HL5X-9 | CGAGTACGAGCGAAGTCTGGTTATACGTCGAGTTGAACGTCGTAACA |
